# Continuous decision to wait for a future reward is guided by fronto-hippocampal anticipatory dynamics

**DOI:** 10.1101/2021.11.16.468916

**Authors:** Reiko Shintaki, Daiki Tanaka, Shinsuke Suzuki, Takaaki Yoshimoto, Norihiro Sadato, Junichi Chikazoe, Koji Jimura

**Author notes:** Correspondence should be addressed to: Koji Jimura, Ph.D. Department of Informatics, Gunma University, 4-2 Aramaki-machi Maebashi, 371-8510, Japan. These authors contributed equally to the study (Reiko Shintaki, Daiki Tanaka). The authors declare no conflict of interest.

## Abstract

Deciding whether to wait for a future reward is crucial for acquiring rewards in an uncertain world and involves anticipating future reward attainment. While seeking for a reward in natural environments, behavioral agents constantly face a trade-off between staying in their current environment or leaving it. It remains unclear, however, how humans make continuous decisions in such situations. Here we show that anticipatory brain activity in the anterior prefrontal cortex (aPFC) and hippocampus underpins continuous stay-leave decision making. Human participants awaited for real liquid rewards available after tens of seconds, and continuous decision was tracked by monitoring dynamic patterns of brain activity. Participants stopped waiting more frequently and sooner after they experienced longer delays and received smaller rewards. When dynamic activity reflecting the anticipation of a future reward was enhanced in the aPFC, participants remained in their current environment, but when this activity diminished, they left the environment for a new one. The anticipatory activity in the aPFC and hippocampus was associated with distinct decision strategies; aPFC activity was enhanced in participants adopting a leave strategy, whereas those remaining stationary showed enhanced activity in the hippocampus. Our results suggest that fronto-hippocampal anticipatory dynamics underlie continuous decision making while anticipating a future reward.

## Introduction

Decision-making in natural environments usually involves multiple factors to be considered. One of the pivotal factors is the time to critical events. It is well known that behavioral agents prefer desirable events that occur sooner rather than later (1, 2).

The devaluation of a temporally remote outcome has been studied in intertemporal decision-making, where future outcomes that vary in both magnitude and time of delivery are evaluated (3, 4). In a standard choice situation, behavioral agents make a one-shot decision between a larger amount of a later reward and a smaller amount of a sooner reward; they then receive the reward after the delay (5–7). However, this decision situation does not allow for the examination of whether behavioral agents are willing to continue to wait or want to abandon waiting after a decision is made. This issue becomes more pronounced when the time to reward delivery is ambiguous (7–12).

A prolonged delay of a desirable event forces behavioral agents to decide whether to continue to wait or to stop waiting. This situation has been studied as a trade-off inherent in delay-of-gratification decisions and foraging behavior (2, 8–11, 13-16). In a typical delay-of-gratification task, behavioral agents wait for a larger amount of reward available later; they can stop waiting for the reward and obtain a smaller amount of reward immediately at any time (2, 8–10, 13). In this context, the ability to persist in waiting for a larger reward is thought to reflect self-control in reward-related decision-making. Similarly, in foraging, behavioral agents constantly face a trade-off between staying at the location of a current food source or exploring to find another to obtain food (7, 11, 14–17). In both contexts, expectations of food acquisition and temporal resources are fundamental factors involved in resolving the trade-off (12, 14, 18).

A common and important valuation in resolving the trade-off is the degree to which behavioral agents anticipate future reward attainment if they remain in their current environment (7, 11, 12, 14, 15). For example, consider an angler who is fishing in a particular spot (Fig. 1A). The angler must constantly decide whether to stay in the current spot or leave it to find another one. The angler will remain if they anticipate that their next catch will be worth the wait; otherwise, they will abandon the spot and look for another. As this example illustrates, the anticipation of future gain plays a key role in stay-leave decision-making (19–22).

**Figure 1.**
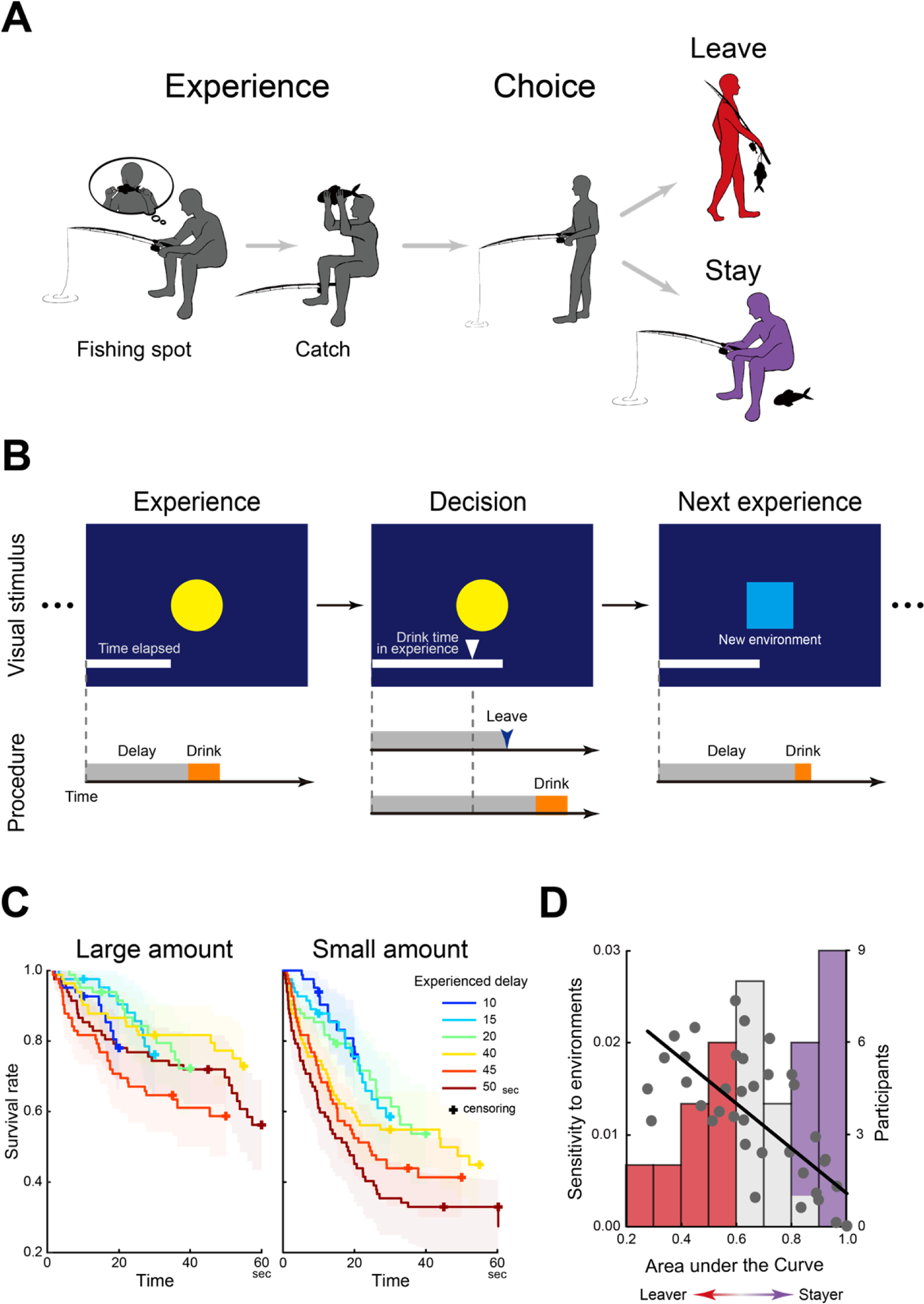
Continuous stay-leave decision-making. A) An illustration of the continuous stay-leave decision. An angler catches a fish, and then decides whether to stay in the current spot or leave in order to find a new one. B) Behavioral procedure. Human participants were given an environment indicated by a picture (yellow circle). In this environment, they first waited for tens of seconds and consumed a real liquid reward (experience trial; *left*). Then, they waited for another liquid reward in the same environment (decision trial; *middle*). During the experience and decision trials, the elapsed time from the start of the trial was indicated by the length of a white horizontal bar extending to the right side of the screen. In the decision trial, the reward delivery time in the experience trial was indicated by a white triangle. In both the experience and decision trials, participants were unsure when they would receive the reward. In the decision trials, however, they were free to leave the current environment at any time and move on to the next experience trial with a new environment. The next environment was indicated by a different picture (blue square; *right*). Note that the picture indicating the environment was unique for each pair of experience and decision trials. C) Survival rates in the decision trial as a function of delay latency. Colors indicate delay durations in the experience trials. *Left*: large reward amount; *right*: small reward amount. Shaded areas indicate 95% CI. Crosses indicate censored events where participants continued to wait until reward delivery. D) The distribution of the area under the curve (AuC) of the survival functions of individual participants. The horizontal axis indicates the AuC of the survival rate, and the frequency of the participants is shown as a bar graph with the vertical axis on the right. Participants were labeled based on the AuC values as leavers (lowest tertile; red) or stayers (highest tertile; purple). The overlying scatter plot shows the correlation between the AuC and the sensitivity to the environmental parameters (reward amount and delay duration). Each plot denotes one participant.

One crucial feature of this behavioral situation is that decisions are made on a continuous-time basis; behavioral agents can stop waiting for a future reward in their current environment at any time. Such stay-leave decision-making has been examined in humans and animals using paradigms and environments optimized for individual species (11, 14, 16–18). In non-human animal studies, choices were made on a continuous basis, which reflects the natural environment and is a characteristic of the aforementioned example (Fig. 1A) (7, 11, 16, 17). In human studies, however, participants did not face this type of continuous trade-off: a stay-leave trade-off was provided each time they received an outcome of a previous choice. Then, they made a one-shot stay-leave choice on a trial-by-trial basis (8, 10, 18, 23).

To address this issue, this study was designed to examine the continuous stay-leave decision-making of humans while awaiting a primary reward. Participants performed a behavioral task for real liquid rewards during functional MRI scanning (Fig. 1B). The current task made it possible to examine the links between anticipation and continuous stay-leave decision-making, and to identify the underlying neural mechanisms. Participants first received a real liquid reward delayed by tens of seconds in a particular environment (experience trial). Then, they waited for another liquid reward in the same environment (decision trial). In this decision trial, they were not exactly sure when they would receive a reward, but could stop waiting and move on to a novel environment whenever they preferred.

## Results

### Stay-leave decisions depend on past experiences

Participants stopped waiting more frequently and sooner after they experienced smaller amounts of a reward and a longer delay, as shown by survival functions reflecting the rate of trials in which participants continued to wait (Fig. 1C; reward amount: χ^2^ = 49.0, P < .001; delay duration: χ^2^ = 26.1, P < .001, log rank test). These results suggest that when the current environment was not promising in terms of acquiring another reward, they abandoned the environment and explored to identify a new one (11, 14, 15, 24, 25). Using laboratory setups where behavioral agents directly experience reward attainment, such continuous stay-leave decision has been observed in a wide range of species from non-mammals (e.g., nematodes, insects, and birds) (16, 17) to mammals (e.g., rodents and primates) (11). Our study is the first to demonstrate that humans make continuous stay-leave decisions for primary rewards in a laboratory setup.

The decision strategy in the decision trial was quantified by the area under the curve (AuC) of the survival function for each trial condition across participants. A smaller AuC reflects the strategy of leaving the current environment for a new one, whereas a larger AuC indicates the strategy of staying in the current environment due to anticipation of a future reward. The AuC was smaller after delivery of a smaller amount of reward [F(1, 40) = 16.6, P < .001] and if there was a longer delay before the reward was delivered [F(1, 40) = 37.9, P < .001]. The adoption of an leave strategy was enhanced by the joint experience of a smaller reward and longer delay [interaction of reward amount and delay duration: F(1, 40) = 11.8, P < .01].

Next, for each participant across trial conditions, the AuC value of the survival function was recalculated, and participants with the highest and lowest AuC tertiles were labeled as stayers (i.e., non-leavers) and leavers, respectively (Fig. 1D). This labeling was intended to characterize individual participants who adopted stay and leave strategies based on the AuC. Importantly, the leavers were more sensitive to environmental factors (reward amount and delay duration) than the stayers (r = −.73, P < .01 with a permutation test; Fig. 1D and Fig. S1), suggesting that leavers more frequently decided to abandon the current environment and move on to a novel one when they received a smaller reward after a longer delay in the experience trial, because the environment was not promising for them.

### Stay-leave decisions involve anticipatory brain activity regarding a future reward

In the decision trial, participants made stay-leave decisions continuously. It is presumed that when they stopped waiting, their anticipation of a future reward was diminished in the current environment. Thus, the survival functions coding for when participants left reflect the temporal characteristics (dynamics) of the reward anticipation (Fig. 1C). We then hypothesized that participants would leave the current environment when brain activity representing the anticipated reward was attenuated. This hypothesis was tested by exploring brain region showing dynamic brain activity reflecting survival functions from the start of the decision trial until the onset of leaving the environment (leave trial; Fig. 2A; Fig. S2A). Exploration of the whole brain revealed that the anterior prefrontal cortex (aPFC) showed dynamic anticipatory brain activity prominently (Fig. 2B; Table S2). The timecourse of aPFC activity showed gradual decrease toward leaving the current environment after its peak (Fig. 2C).

**Figure 2.**
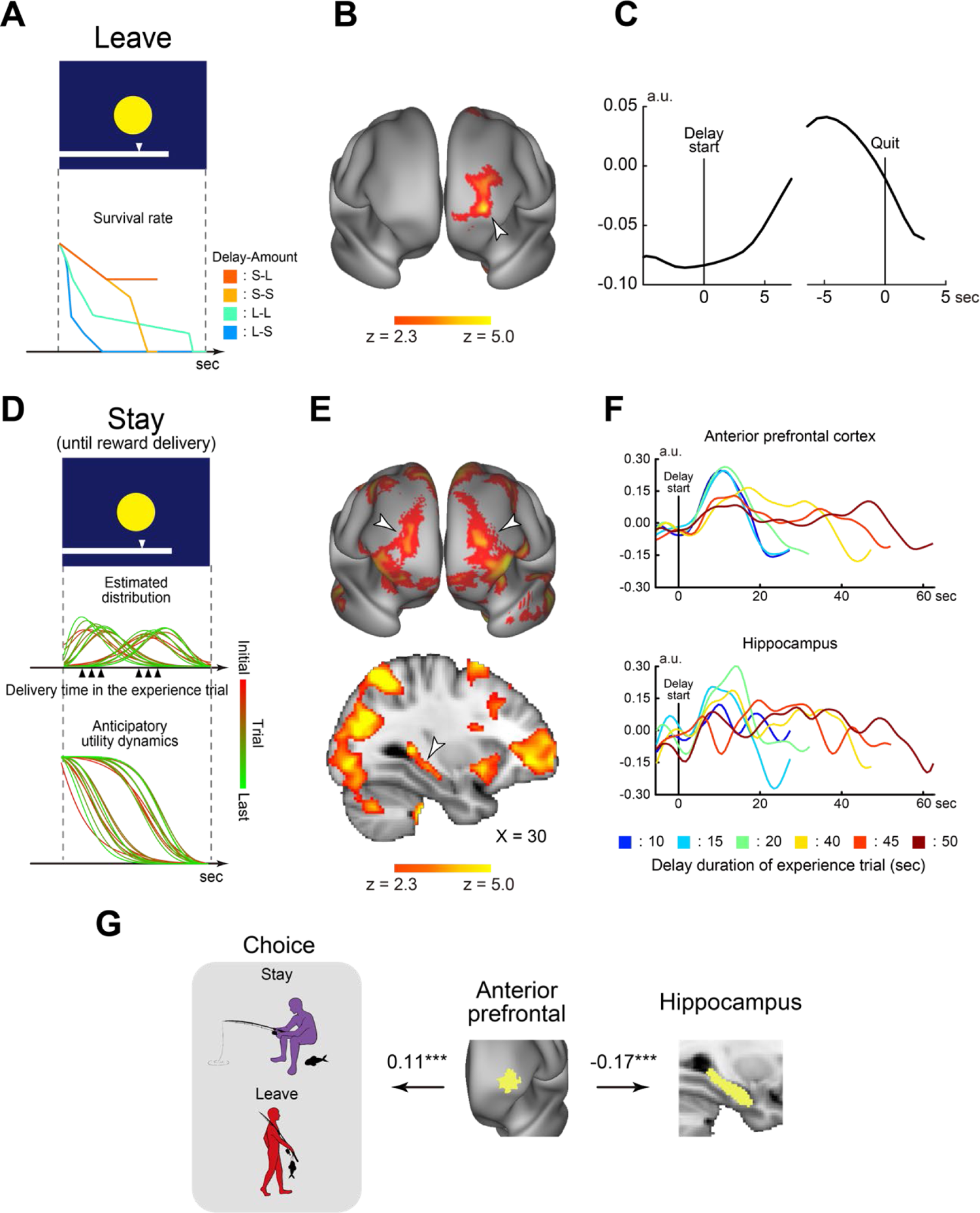
The anterior prefrontal cortex (aPFC) and hippocampus (HPC) showed anticipatory activity during continuous stay-leave decision-making. A) Anticipatory dynamics during the leave trials were modeled by an individual’s survival function for each condition. Delay: Short (S)/Long (L); Reward amount: Small (S)/Large (L). B) Brain regions showing activation dynamics of the survival function during the delay period in the leave trials. Maps are overlaid onto the three-dimensional brain surface. The color bar indicates significance levels. The white arrowhead indicates the aPFC. C) The timecourse of brain activity in the aPFC. D) The probability density distribution of reward delivery expectation was updated throughout the experience and decision trials (*middle*). Anticipatory utility (AU) dynamics during the stay trials were modeled as inverse dynamics of the cumulative probability of the density distribution (*bottom*). The colors of the lines indicate the trials in the color bar on the right. E) Brain regions showing AU dynamics during the delay period of the stay trials. Anterior view (*top*); sagittal section (*bottom*; X = 30). The arrowheads indicate the aPFC (*top*) and HPC (*bottom*). The formats are similar to those in Fig. 2B. F) The timecourses of activity in the aPFC (*top*) and HPC (*bottom*). G) Choice behavior in the decision trials was predicted by the aPFC activity, and the HPC activity was negatively coupled with the aPFC activity. Yellow areas indicate ROIs in the aPFC, defined based on prior studies, and in the HPC, which was defined anatomically. Values indicate regression coefficients. ***: P < .001.

We next examined anticipatory brain activity in trials where participants continued to wait until a reward was delivered (stay trial). In theory, the survival function is not suitable to examine anticipatory activity in the stay trials because this function encodes when to stop waiting. Alternatively, the dynamics of anticipatory activity were modeled based on the theory of anticipatory utility (AU), based on the assumption that anticipation of a future reward itself provides pleasure and confers current utility (6, 20, 26). Specifically, AU was modeled based on participants’ expectation of reward delivery formulated by a probability density function, which was updated every time after participants performed decision trial (22) (Fig. 2D; see Methods and Fig. S2B/C).

Whole-brain exploratory analysis revealed a strong AU effect in the aPFC (Fig. 2E *top*). Interestingly, this region also showed anticipatory activity in the leave trials (Fig. 2B). The timecourse of aPFC activity revealed the temporal dynamics of AU (Fig. 2F *top*). The hippocampus (HPC) also showed a strong AU effect (Figs. 2E/F *bottom*; Table S3), though such anticipatory activity was absent in the leave trials (Fig. S3). The HPC activity during the stay trials may reflect reproduction of the pleasure of anticipation encoded while participants waited for a reward in the experience trial (27–33), which may not be the case in the leave trials.

As suggested by previous studies, the value of a future reward was increased monotonically while the reward was anticipated (5, 34). The dynamics of the reward value are complementary to AU dynamics (6, 10, 21, 22), and were modeled as the upcoming future reward (UFR; Fig. S2D). The UFR effect was observed in multiple frontal and parietal regions (Fig. S4A), but medially to the brain regions showing the AU effect (Figs. 2B/E), consistent with previous studies (6, 22).

To compare the fit of AU and survival functions to the fMRI data, we performed a supplementary analysis by reversing the relationships between trials and models. Specifically, activity dynamics were modeled by the AU model for the leave trial, and by the survival model for the stay trial. In the aPFC, both the AU and survival models showed a significant effect (ts > 3.4; Ps < .005; Fig. S5), but the survival model better fit to the leave trials than the stay trials [t(35) = 2.5; P < .05]. This suggests that aPFC activity is linked to leaving the current environment because the survival model directly codes the probability of leaving. In the HPC, on the other hand, in both models the effects were significant in the stay trials (ts > 2.1; Ps < .05) but were absent in the leave trials (ts < 0.6; Ps > .6), suggesting that HPC activity leads to remaining in the current environment.

### Stay-leave decisions are governed by anticipatory aPFC activity

Our results demonstrated that 1) the aPFC and HPC showed anticipatory activity in the stay trials (Fig. 2E), and 2) leaving an environment was associated with attenuated aPFC activity (Fig. 2C). We then hypothesized that 1) the continuous decisions would be predicted by aPFC activity, and 2) the HPC would functionally coordinated with the aPFC when the environment was not abandoned before reward delivery.

To test the former hypothesis, we performed a regression analysis in which anticipatory activity predicted whether there was departure until reward delivery or when a new environment was sought out during the decision trials. The analysis was implemented by a multi-level mixed-effects general linear model (GLM) involving the aPFC or HPC activity in individual trials as predictors. The aPFC activity successfully predicted the choice behavior (*Z* = 3.8; P < .001; Fig. 2G), indicating that weaker aPFC activity is associated with leaving the current environment. The HPC did not predict the choice behavior (*Z* = 0.77; P > .44).

The latter hypothesis was then tested by examining the functional connectivity between the HPC and aPFC based on a mixed-effects GLM. aPFC activity was negatively coupled with HPC activity in individual trials (*z* = 3.4; P < .001), suggesting that the aPFC and HPC play complementary roles in the choice to remain in the environment during the continuous stay-leave decision-making.

### aPFC and HPC differentially code decision strategies

The degree of adoption of the leave and stay strategies is reflected in individual participants’ AuC value of the survival functions (Fig. 1D) and the anticipatory activity dynamics were observed in the aPFC and HPC in the decision trials (Figs. 2B-F). Then, we next asked whether participants with distinct stay and leave strategies (i.e., stayer and leavers) show differential involvement of these brain regions. To test this possibility, we explored brain regions showing cross-subject correlations between AU effects in the stay trials and the AuC values of individuals’ survival functions.

The aPFC showed a strong negative correlation between the AU effect and AuC, (Fig. 3A *left*; Table S4) indicating that leavers showed strong AU effects. In contrast, the HPC showed a positive correlation (Fig. 3B *left*; Table S4), indicating that the stayers showed even stronger AU effects. Scatter plots of the AU effect against AuCs showed clear negative and positive correlations in the aPFC and HPC, respectively (Figs. 3A/B *middle*). Notably, these regions also showed strong AU effects (Figs. 2B/E). To show this correlation more specifically, timecourses of MRI signals in these regions were extracted for stayers and leavers as defined in Fig. 1D. Leavers showed strong anticipatory activity in the aPFC but not in the HPC, whereas stayer showed strong anticipatory activity in the HPC but not in the aPFC (Fig. 3A/B *right*). These results demonstrate that distinct decision strategies involve the aPFC and HPC differentially in the stay trials.

**Figure 3.**
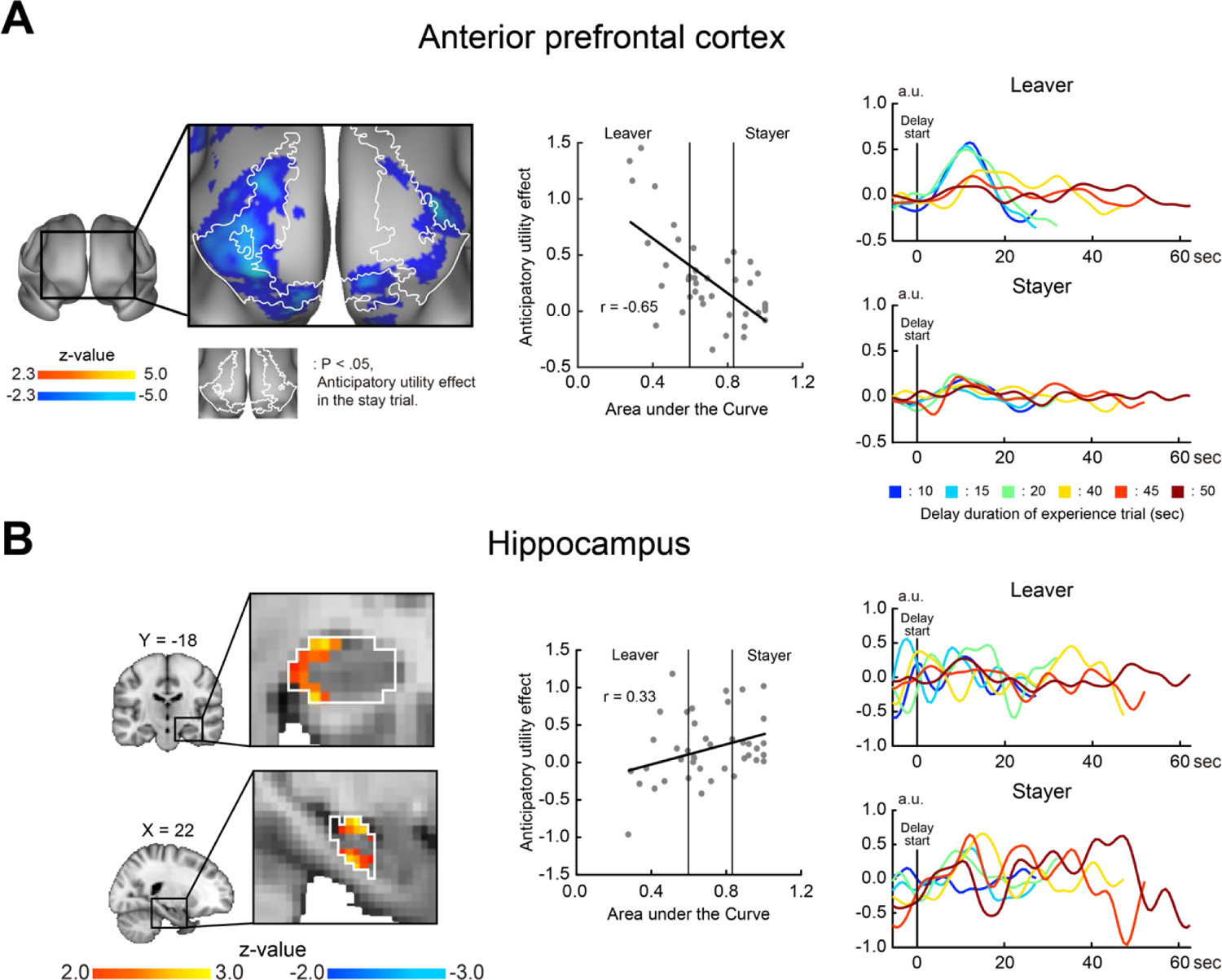
Stayers and leavers exhibit differential involvement of the aPFC and HPC. A) Statistical maps of correlations between AU effects and AuCs were overlaid onto a three-dimensional brain surface (*left*). Cool and hot colors represent greater AU effects in leavers (smaller AuCs) and stayers (greater AuCs), respectively. The color bars indicate significance levels. The black rectangle on the map (*left*) represents the area magnified on the right. White closed lines indicate the aPFC regions showing AU effects in Fig. 2E *top*. *Middle*: A scatter diagram of AuC values and AU effects in the aPFC. Each plot denotes one participant. Timecourses of brain activity in the stay trials for leavers (*right top*) and stayers (*right bottom*). B) Statistical correlation maps in the HPC (*left*). White closed lines indicate anatomical borders of the HPC. Formats are similar to those in Fig. 3A.

### The HPC encodes anticipatory experience in a novel environment

Before a decision trial, participants received a delayed reward in an experience trial with a novel environment. We then asked whether anticipatory brain activity in the experience trials was associated with choice behavior in the decision trials. We explored anticipatory brain activity in the experience trials by modeling AU and UFR dynamics (Fig. 4A; Fig. S6A), as in the stay trials (Fig. 2D).

**Figure 4.**
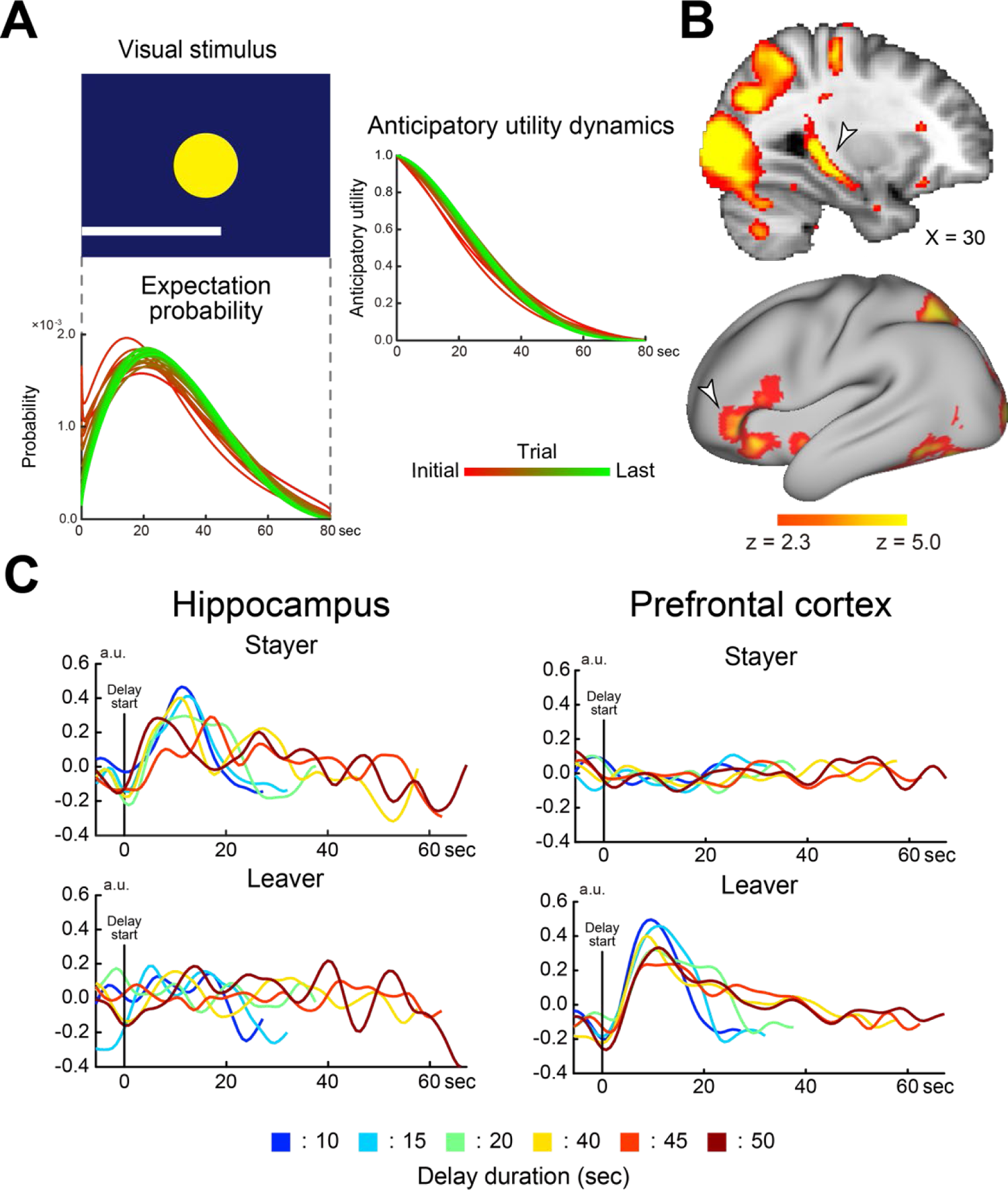
The experience of anticipation is encoded in the HPC. A) The AU dynamics in the experience trials (*right*) were modeled similarly to those in the stay decision trials. Formats are similar to those in Fig. 2D. B) Brain regions showing AU dynamics during the delay period of the experience trials. The arrowheads indicate the effects in the HPC (*top*) and aPFC (*bottom*). Formats are similar to those in Fig. 2E. C) Timecourses of brain activity during the delay period. Timecourses are shown for the HPC (*left*), aPFC (*right*), stayers (*top*), and leavers (*bottom*).

Whole-brain exploratory analysis revealed that the HPC and ventrolateral prefrontal cortex (vlPFC) showed a strong AU effect (Fig. 4B; Fig. S6B/C). The vlPFC region was spatially segregated from the aPFC region that showed an AU effect in the decision trial (23.7 mm apart).

Importantly, the stayers showed a stronger AU effect in the HPC, whereas in the leavers this effect was stronger in the vlPFC (Fig. 4C; Fig. S6D). It is interesting that in the decision trials, a stronger AU effect in the HPC was also observed in the stayers, suggesting that the stay strategy may be guided by HPC-dependent encoding and reproduction of anticipation of a future reward (35–37).

## Discussion

In the current study, humans made continuous stay-leave decisions regarding real liquid rewards available after tens of seconds, and the decisions were tracked by temporal dynamics of brain activity reflecting anticipation of a future reward. When the anticipatory activity in the aPFC diminished, they abandoned the current environment. The hippocampal activity was enhanced in participants remaining stationary, whereas those adopting a leave strategy showed strong aPFC activity. As such, anticipatory dynamics in the fronto-hippocampal mechanisms underlie continuous stay-leave decision-making about a future reward.

Our study highlights the distinctive roles of the aPFC and HPC during reward-seeking behavior in temporally uncertain environments (Fig. 5). Specifically, the aPFC encodes choice behavior and is associated with leave strategy in the decision trials, but is not engaged in the experience trials, suggesting that the aPFC is sensitive to the current pleasure (utility) from moment to moment in the decision trials (6, 19, 22, 38). On the other hand, the HPC is involved in the experience trials and is associated with the stay strategy, but it is unrelated to leaving the current environment. The greater HPC effect in the stayers may reflect enhanced reproduction of the experience of reward anticipation (35, 36, 39–41), as suggested by prior non-human mammal studies demonstrating that the HPC encodes and reproduces temporal series of events (27–33).

**Figure 5.**
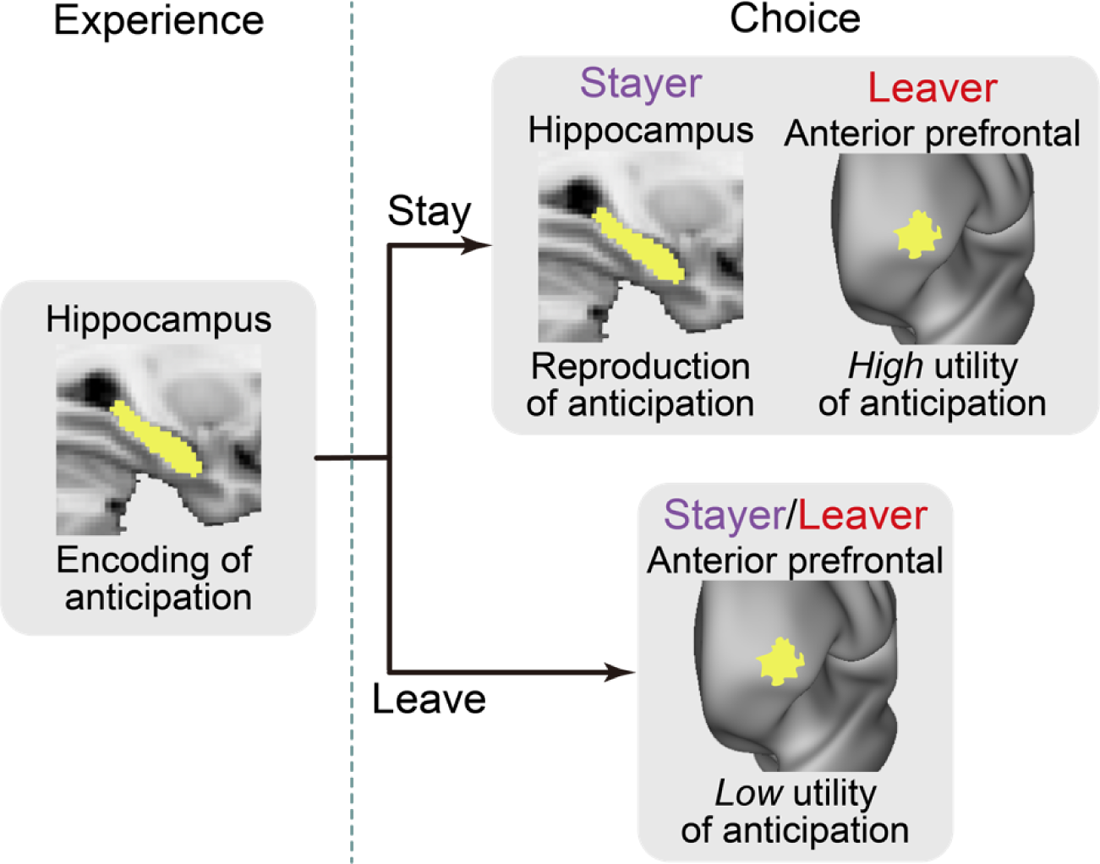
Schematic diagram of human prefrontal-hippocampal mechanisms underlying reward-seeking behavior in temporally uncertain environments.

The current aPFC region is located in a polar region in the prefrontal cortex within Brodmann area 10, which is disproportionately developed in humans (42). This region shows dynamic AU representation when a delayed reward is anticipated (6, 22). The region is also related to thinking about both future events (19, 39, 43, 44) and exploration of a novel environment involving anticipatory evaluation of the current and novel environments (21). Thus, our results highlight that a prefrontal region unique to humans represents a desirable future event while anticipating a future reward.

The current finding that gradual decrease of aPFC anticipatory activity toward leaving the current environment (Fig. 2C) contrasts with previous rodent, primate, and human studies showing that increased neuronal activity in the medial frontal area triggers exploration of a new environment (14, 45) and predicts when the decision to leave the environment is made (46–51). In *C. elegans*, on the other hand, decreased activity of serotonergic neurons drives exploration (17), which better fits our finding in terms of activity reduction associated with leaving the current environment. It has also been demonstrated that stay-leave strategies during foraging of *C. elegans* are affected by genetic variations in adrenergic receptors (16) and serotonergic neuronal activity (17). Our findings of individual differences in stay-leave strategies (Figs. 3A/B) and the underlying prefrontal-hippocampal mechanisms in humans may be reflected in molecular neurobiological mechanisms shared across species.

The continuous decision required during the decision trials involves a trade-off between staying in the current environment and leaving to locate a novel environment. A similar trade-off is a choice between exploration and exploitation, where behavioral agents repeatedly make choices among options to maximize long-term reward attainment (bandit task). In such situations, the agent’s valuation of options was updated based on their history of choice outcomes (23, 52, 53). Interestingly, human studies using abstract rewards in hypothetical situations showed that exploration of an uncertain option involved a region in the anterior prefrontal cortex (23). The current study suggests that the involvement of the aPFC reflects stronger anticipation for another choice.

It has been argued that given the complexity of the real world, foraging behavior is governed by several environmental factors (54). Such factors include success and failure of prey acquisition (45), duration since previous prey acquisitions, costs of traveling between feeding sites (14), cost of engagement in the current site (18), and resources available in the environments (55). These factors are well explained by the marginal value theorem and other formal models (24). In the real world, predators, competitors, and nutrition can also be important (15). In the current study, we aimed to examine the effect of anticipating future reward attainment, and then manipulated the amount of liquid rewards and the duration to the reward outcome. Consideration of exploration costs of humas is associated with the anterior cingulate cortex (18). By contrast, we focused on anticipation of a future reward, and did not observe involvements in the anterior cingulate cortex.

In foraging situations, behavioral agents attempt to maximize long-term prey acquisition based on continuous consideration of the stay-leave trade-off, like in the decision trials in the current study. The maximization of future reward attainment has been examined in decision-making situations involving a choice between a larger delayed reward and a smaller reward available sooner, formulated as intertemporal choice (6, 7, 20, 22). The preference in intertemporal choice reflects impulsivity and self-control in reward-seeking behavior (5). Importantly, in intertemporal choice, the optimal choice to maximize long-term reward attainment, i.e., by choosing the larger delayed reward, is always obvious to the agents. By contrast, in foraging situations, the optimal choice is not always obvious to agents because there is uncertainty about future prey acquisition in the surrounding environment. Thus, the continuous stay-leave choice in the current decision trials may not be compatible with the impulsivity-self-control spectrum in intertemporal choice, and choice preference may be associated with other trait constructs reflecting anticipation of future desirable events (20, 26).

In the literature on foraging, there are discrepancies between human and animal studies, despite the ethological generality and importance of foraging among a wide range of species. In non-human animal studies, behavioral agents directly foraged for primary rewards (7, 11, 14, 16, 17, 45), whereas previous human studies used secondary rewards mainly in hypothetical situations (18, 50, 55) (cf. entertaining videos as rewards in (11)). Due to these discrepancies, it was challenging to compare foraging and underlying biological mechanisms across species. The current study bridged the gap between human and animal studies, providing important consistency across species.

Variability in what behavioral agents anticipate is another important issue to be controlled. In non-human animals, stay-leave decision-making has been examined in real situations using primary rewards involving nutrition (e.g., food or liquid) (11, 14, 16, 17). In humans, on the other hand, most paradigms have used secondary rewards (e.g., money or tokens) in hypothetical situations (8, 10, 18). However, humans make choices differently for real primary rewards and hypothetical secondary rewards (56). These discrepancies in types of choices and rewards type make it hard to compare stay-leave decision-making across species. Thus, the use of real nutritious rewards under a continuous trade-off in a human experiment should provide significant insight to extend our understanding of stay-leave decision-making.

## Acknowledgments

This work was supported by JSPS Kakenhi (21H05060, 20K07727, 17K01989, and 26350986 to K.J.); NIPS Cooperative Study Program (20-639, 19-635, 18-633, 17-6229, and 21-544 to K.J.); ABiS (16A-073-M02 to K.J.); AMED (JP21dm0207086 to J.C.) We thank Drs. Kotaro Oka, Teppei Matsui, and Kentaro Miyamoto for their scientific comments on the manuscript. We thank Mimu Yabuta for illustration assistance.

## Author contributions

D.T., N.S., and K.J. designed research; R.S., D.T., T.Y., J.C., and K.J. performed research; R.S., D.T., S.S. and K.J. contributed analytic tools; R.S., D.T. and K.J. analyzed data; R.S. and K.J. wrote the first draft of the paper; R.S., S.S., N.S., J.C., and K.J. edited the paper; R.S. and K.J. wrote the paper.

## Materials and Methods

### Participants

Human participants (N = 41; age range, 20-28 years; 20 females) were right handed and had no history of psychiatric or neurological disorders. Written informed consent was obtained from all participants. All experimental procedures were approved by the institutional review boards of Keio University and the National Institute of Physiological Sciences. Participants were instructed not to drink any liquid for 4 hours before the experiment and received 8,000 yen for participation. Five participants did not leave the current environment in any decision trial, and these participants were excluded from the analyses examining trials where participants stopped waiting.

### Reward

The current study used commercially available drinks as a reward that could be immediately consumed. Before the experiment, participants were provided with a list of the following beverages: apple, orange, grape, grapefruit, lychee, pear, and mixed fruit juices; probiotic drinks; barley tea; and water. Each participant was then asked to choose the drink that would serve as their reward.

### Apparatus

E-prime programs (Psychology Software Tools) controlled the task as well as the delivery of liquid rewards via a syringe pump (SP210iw; World Precision Instruments). Liquids from two 60-ml plastic syringes mounted on the pump were merged into one tube and then delivered to the participant’s mouth through a silicon tube (Movie S1).

The flow rate of each syringe was set to 0.75 ml/s, and thus the reward flowed continuously at a rate of 1.5 ml/s. Participants were able to control the liquid flow. Reward delivery continued as long as they pressed a button on a box that they held in their right hand; delivery paused if they released the button, and resumed when they pressed the button again.

### Behavioral procedures

During functional MRI scanning, human participants performed a behavioral task seeking for real liquid rewards delayed by tens of seconds (Fig. 1B; Movie S1).

Participants first experienced a delayed reward in a novel environment indicated by a picture presented on the center of the screen (experience trial), and then waited for another reward in the same environment (decision trial). Importantly, in the decision trial, participants were unsure when they would receive the reward; however, they were able to stop waiting and move on to a novel environment whenever they preferred. They alternately performed the experience and decision trials, and an environment was used only for one pair of the experience and decision trials. Each environment was indicated by a unique picture.

Prior to each experience trial, participants were presented with a visual message to wait until delivery of a reward, and to press the button to start the trial. The visual message was presented until the participants pressed the button with their right thumb, at which point the visual message disappeared and a fixation cross was presented for 3 s. Then, a picture indicating an environment was presented, and the delay period started. During the delay period, the elapsed time from the start of the trial was indicated by a white horizontal bar that extended from the left to the right side of the screen every 250 ms. A full bar extending to the right end of the screen corresponded to 80 s, which was longer than the maximum delay duration in the decision and experience trials (60 s). At the end of the delay, a visual message was presented indicating that the reward was ready, and participants consumed the liquid reward. After they consumed all of the liquid, a fixation cross was presented for 13 s.

Then, participants waited for another reward in the same environment (decision trial). Before starting of the decision trials, they were presented with a visual message telling them to wait for another delayed reward, to stop waiting by pressing the button at any time, and to press the button to start the trial. When they pressed the button, the visual message disappeared and a fixation cross was presented for 3 s. Then, the picture indicating the environment in the experience trial was presented again, and participants started waiting. While they were waiting, the elapsed time was indicated by a white elongating bar as in the experience trials. Additionally, a white triangle was shown above the white bar, indicating the reward delivery time in the experience trial.

If participants waited until the delivery of a reward (stay trial), a visual message indicating that the reward was ready was presented, and participants consumed the liquid reward. If they stopped waiting before the reward delivery (leave trial), the environmental information (center picture, white bar, and triangle) disappeared, and a fixation cross was presented. After an inter-trial interval (ITI), the next experience trial started with a new environment. The duration from the onset of the decision trial to the end of the ITI was 75-165 s, which varied by reward amount and delay duration (see below), but was independent of participants’ choices in the decision trials. This meant that the ITI was longer in the leave trials, but no participants reported that they noticed a longer ITI when they stopped waiting. This ensured that participants did not consider the duration of the ITI when making decisions during the decision trials.

Participants performed a total of 24 pairs of experience and decision trials during fMRI scanning, and each scanning run consisted of four pairs of the trials. Table S1 lists the delay durations of all trials. Specifically, there were six delay durations in the experience trials (10, 15, 20, 40, 45, and 50 s). The short-delay condition comprised of the 10-, 15-, and 20-s delays, and the long-delay condition consisted of the 40-, 45-, and 50-s delays. In the decision trials, the delay duration was shorter or longer than that of the experience trials (Table S1). The amount of the delayed reward was either 3 ml (small amount) or 12 ml (large amount). Thus, the current experiment was based on a 2 x 2 factorial design (delay duration: short/long; reward amount: large/small). These conditions appeared pseudorandomly within one scanning run and across scanning runs.

Before fMRI scanning, participants received instructions for the task using a computer display. Participants were told that reward delivery times in decision trials could be shorter or longer than those in experience trials (the latter indicated by a white triangle). To familiarize participants with the task, two pairs of experience and decision trials were performed as practice trials in the MRI scanner [(delay duration in the experience trial, reward amount, delay duration in the decision trial): (5 s, 6 ml, 15 s), (25 s, 6 ml, 15 s)]. The total liquid consumption in an entire experimental session ranged from 192 ml to 384 ml.

Each environment consisted of one of 24 unique pictures (six shapes x four colors). One color was used only once in each scanning run. Participants were told that the picture used in one trial pair was unrelated to that used in other trial pairs. Pictures used in practice trials were not used in scanning trial pairs.

### Survival rate analysis and assessment of stay-leave strategies

To assess participants’ behavior in the decision trials, we used survival analysis based on the Kaplan-Meier method (57). This analysis examined how long participants remained in an environment by estimating the frequency with which they continued to wait until time *t* during the delay period of the decision trials (survival rate). Specifically, the rates were calculated based on observations of leaving the environment (event occurrence) and completion of the delay period of the decision trial (censoring). The survival rate was formulated as

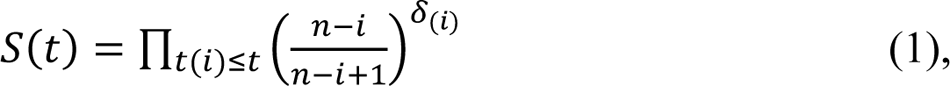

where *S*(*t*) represents the estimation of the survival rate at the time *t*, *n* is the sample size, and *t*_(*i*)_ denotes the time of the *i*th event occurrence. δ_(*i*)_ is an indicator function that takes a value of 0 if the *i*th observed event is a censoring observation and 1 if it is a non-censoring observation. The survival rate was calculated for each reward amount and delay duration of the experience trials across participants using *ecdf* implemented in MATLAB ver. 2017a.

The stay-leave strategy in the decision trial was quantified by the area under the curve (AuC) of the survival function for each trial condition across participants. A smaller AuC reflects the strategy to leave the current environment for a new one, whereas a greater AuC reflects the stay strategy in the current environment due to anticipation of a future reward. The AuC values were calculated for each delay and amount condition and compared across trial conditions.

The AuC values were also calculated for each participant across conditions to quantify individuals’ choice strategies in the decision trials. Individuals with greater AuCs waited longer (i.e., adopting a stay strategy) whereas those with smaller AuCs waited for less time (i.e., adopting a leave strategy). Given this characterization of the individuals’ AuCs, we classified participants into three groups based on AuC values, and labeled the highest tertile as stayers and the lowest tertile as leavers.

### Sensitivity to environmental condition

To examine how environmental parameters (reward amount and delay duration in the experience trials) affected choice behaviors during decision trials, we quantified individuals’ environmental sensitivity. We first hypothesized that the duration that participants would wait in a decision trial would be predicted by the reward amount and delay duration in the experience trials. This hypothesis was formulated by the following regression model for each participant:

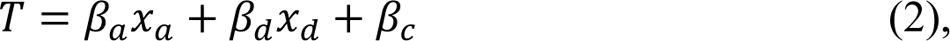

where *T* is (waiting duration) / (time until reward delivery during decision trials) (= 1, if participants waited until reward delivery); *x*_a_ and *x*_d_ are the reward amount and delay duration, respectively; β_a_ and β_d_ are the regression coefficients for the reward amount and delay duration, respectively; and β_c_ is a constant term.

Data for the estimated coefficients of β_a_ and β_d_ were collected from all participants and averaged across participants. The significance of the averaged β_a_ and β_d_ was tested by permutation tests. Specifically, within participants, the parameters of reward amount and delay duration were randomly shuffled and relabeled, and then the same regression analysis was performed and coefficients were collected and averaged across participants. Crucially, this random relabeling was performed within participants so that participants’ individuality was preserved. This procedure was repeated 5000 times. Then, group-level averages from 5000 randomizations were collected, which provided for null distributions of β_a_ and β_d_ . β_a_ and β_d_ were 0.0056 ± 0.0075 (mean ± SD) (P < .001) and −0.0072 ± 0.0059 (P < .001), respectively, suggesting that participants waited longer if they received a larger reward and/or if they received a reward after a shorter delay. Thus, a more positive β_a_ indicates that the wait duration was longer when a participant received a larger reward. Likewise, a more negative β_d_ indicates that the wait duration was longer when a participant received a reward after shorter delay.

Then, β_a_, β_d_, and β_c_ were estimated for each participant. The sensitivity to environmental parameters *se* was defined as

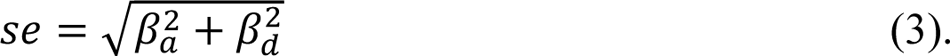

To test whether the *se* value of individual participants was associated with their stay-leave strategy in the decision trials, a correlation coefficient was calculated between the *se* values and the AuC values of the survival function across participants. Notably, AuC represents individuals’ overall choice strategies across trials, whereas *se* represents trial-by-trial variability in choice behavior within individuals. Thus, *se* and AuC represent conceptually independent constructs. However, they were not quantitatively independent, because both AuC and *se* were calculated based on the behavioral data in the decision trials. Due to the quantitative dependence of these values, they could be pseudo-correlated, which requires careful consideration when calculating a null distribution of the correlation. Because the pseudo-correlation is derived from the individuality of the behavioral data, when calculating a null distribution, we preserved the individuality by permuting experimental conditions within individuals, similarly to the testing for the mean β_a_ and β_d_ values across participants as stated above. As expected, the null distribution shows a shift toward the negative direction (Fig. S1). Then, significance of the correlation was tested based on the null distribution.

### Imaging procedure

fMRI scanning was conducted on a whole-body 3T MRI system (Siemens Verio, Germany). Functional images were acquired using multi-band accelerated gradient-echo echo-planar imaging [repetition time (TR) = 800 ms; echo time (TE) = 30 ms; flip angle (FA) = 45 degrees; slice thickness, 2 mm; in-plane resolution, 2 x 2 mm; multi-band factor (MBF) = 8; 80 slices]. Whole brain scanning with high temporal resolution allowed us to perform exploratory analyses of temporal dynamics during the delay period with sufficient scanning frames. Each run involved 635 volume acquisitions (508 s), and six functional runs were performed (total 3990 volumes). The initial 10 volumes of each run were excluded from imaging analysis to take into account the equilibrium of longitudinal magnetization. High-resolution anatomical images were acquired using an MP-RAGE T1-weighted sequence (TR = 2500 ms; TE = 4.32 ms; FA: 8 deg; 208 slices; slice thickness, 0.8 mm; in-plane resolution, 0.8 x 0.8 mm^2^).

### Imaging analysis procedures. Preprocessing

Functional images were preprocessed using SPM12 (http://www.fil.ion.ucl.ac.uk/spm/). All functional images were first temporally aligned across the brain volume, corrected for movement using correction for rigid-body rotation and translation correction, and then registered to the participant’s anatomical images to correct for movement between the anatomical and function scans. Participants’ anatomical images were transformed into a standardized MNI template. The functional images were then registered to the reference brain using the alignment parameters derived for the anatomical scans. The data were resampled into 2-mm isotropic voxels, and spatially smoothed with a 6-mm full-width at half-maximum Gaussian kernel.

In order to minimize motion-derived artifacts due to consumption of liquid rewards (6, 22, 38), functional images were further preprocessed by general linear model (GLM) estimations with motion parameters and MRI signal time courses (cerebrospinal fluid, white matter, and whole brain), and their derivatives and quadratics as nuisance regressors (22, 58–60) based on *fsl_regfilt* implemented in the FSL suite (http://www.fmrib.ox.ac.uk/; ver. 5.0.9). The residual of the nuisance GLM was used for standard GLM estimations to extract event-related brain activity as described below.

The mean magnitudes of absolute head motion were less than 0.23 mm for translation and 3.8 x 10^-3^ radian for rotation, which is consistent with our prior studies (6, 22, 38) (see (38) for details). Critical image distortions or signal drop-outs were not observed, but robust brain activity was extracted in the primary motor and gustatory cortices during liquid consumptions, which assured us that as in our prior studies (6, 22, 38), motion-derived artifacts did not critically contaminate the current analyses.

### General linear model

#### Single level analysis

A GLM approach was used to estimate trial event effects of brain activity. Parameter estimates were performed by *feat* implemented in the FSL suite. Events of particular interest were defined as periods during which participants waited for a future reward, i.e., the delay periods of the decision and experience trials. The current analysis focused on the temporal dynamics of brain activity during these events (6, 22).

#### Decision trial: leave before reward delivery

During the decision trials, participants anticipated a future reward and continuously decided whether to stay in or leave the current environment. When the anticipation of a future reward diminished, they left the current environment. Thus, we hypothesized that this departure occurred when brain activity representing the anticipation of a future reward in the current environment was attenuated. To model temporal changes in anticipatory activity, we used the survival function of the decision trials because continuous decision and anticipation are reflected in temporal changes in survival rates. This hypothesis was then tested by exploring brain regions showing dynamic brain activity associated with the survival functions.

Specifically, the anticipatory dynamics of brain activity were modeled using the survival functions calculated in behavioral analysis (Fig. 2A). For each participant, we estimated survival functions for each of the four trial conditions (reward amount: large/small; delay duration: short/long; see Table S1). Then, for each leave trial, the delay period was coded by the survival function from the onset of the decision trial until the button press indicating that the participant was leaving the current environment.

Alternative temporal dynamics were modeled by the inverse of the survival function (1 - survival function = death function), which is simultaneously coded in the GLM. Then, the survival and death models were convolved with a canonical hemodynamic response function (HRF) (Fig. S2A).

By definition, the survival and death functions are anticorrelated, but when convolved with the HRF, they produced dissociable BOLD regression functions (Figs. S2A/4B). Details regarding the orthogonality of the regressors and the control analyses for inspecting multicollinearity are described in the section on GLM estimations and control analyses below.

### Decision trial: stay until reward delivery

We examined anticipatory dynamics of brain activity during the delay period of the decision trial where participants stayed in an environment until reward delivery. The survival function of the decision trials is not suitable for modeling these dynamics because the survival function reflects when participants stopped waiting, whereas in the stay trials, they continued waiting until reward delivery.

One useful theorization of the anticipation of desirable events occurring in the future is the current utility of anticipation (anticipatory utility: AU) reflecting the pleasure of future anticipation (26). In accordance with AU, our previous studies modeled brain activity dynamics while participants were awaiting a future reward (6, 22). Notably, in the current decision trials, participants waiting for a reward were unsure about when this reward would be delivered. In our previous study (22), AU in these uncertain situations was modeled based on expectation of future reward delivery, which was updated following every experience of a delayed reward using a Bayesian learning inference approach (52). In the current study, this framework was applied to modeling the anticipation of a future reward in the stay trials.

In the decision trials, although participants were unsure when the reward would be delivered, it was possible to estimate when a reward would become available based on the previous reward experience in the same environment (i.e., the experience trial immediately before the decision trial). The expectation should also depend on past decision trial experiences where participants waited until rewards were delivered or stopped waiting before delivery. Thus, the current modelling assumed that prior to the decision trial, participants approximated the delivery time based on the immediately preceding experience trial, and the approximation was updated every time after participants performed a decision trial.

The current analysis modeled the expectation of reward delivery and its updates using a Bayesian inference learning approach. We assumed that the expectation was expressed by a probability density distribution of the reward outcome, which was formulated based on a beta distribution as a function of the expected delivery time. A beta distribution was used because of its finite ends, which is consistent with participants’ understanding of the maximum possible duration in the decision trials. It should be noted that the amount of time that had passed in an individual decision trial was shown by the length of the white bar, which did not extend beyond the right end of the screen (corresponding to 80 s; see Behavioral procedures).

The updates were implemented differently depending on whether participants waited until the reward delivery (stay) or left before it was delivered (leave) in the previous environment. For the stay trials, the update was based on 1) the reward delivery time in the immediately preceding experience trial, and 2) the disparity of reward delivery times between the experience and decision trials in the previous environment (Fig. 2D and Fig. S2B *top*). Specifically, Bayesian inference was performed as

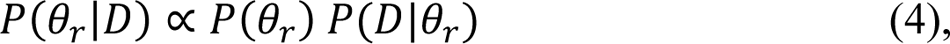

where *D* represents the delivery time of the decision trial and θ_r_ is a parameter that determines the probability density defined by 1) the delivery time of the experience trial indicated by the white triangle and 2) the disparity between the triangle and the right end of the white bar when the reward was delivered.

For the leave trials, on the other hand, the update was based on 1) the reward delivery time in the immediately preceding experience trial, as in the stay trials, but also on 2) the time when participants stopped waiting (Fig. S2B *bottom*). Specifically, Bayesian inference was performed as

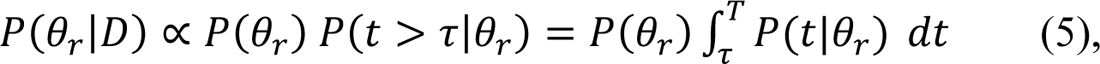

where τ and *T* indicate the time participants left the current environment and the maximum possible delay (80 s), respectively. This formulation reflects the fact that the reward would have been delivered between times τ and *T* because leaving at time τ indicates that the reward had not yet been delivered by this time.

Then, the probability density function (*PDF*) in the stay and leave trials was calculated as

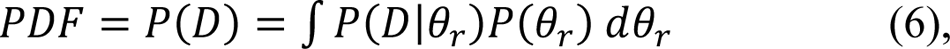

where *P*(*D*|θ_r_) takes a beta distribution. The average of the initial distribution was 15 s (i.e., average of delay durations in the practice decision trials).

To model anticipatory utility dynamics during the decision trials, the PDF was first integrated from the start of the delay to time *t*, defined as cumulative probability (*CP*),

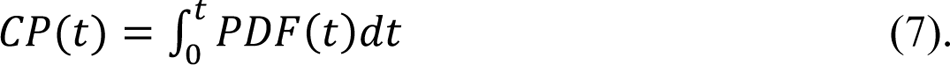

The idea of this integration is that as the delay period elapsed, participants’ expectations of receiving a reward continued to increase, although they did not exactly know when the reward would become available. *CP*(*t*) is similar to a hazard rate in that it reflects an estimation of the probability that the reward is likely to become available(10, 61). As time *t* approaches its maximum value (i.e., 80 s), *CP*(*t*) reaches the upper limit,

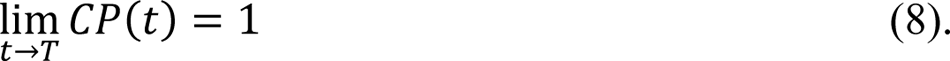

Then, *AU* was defined as

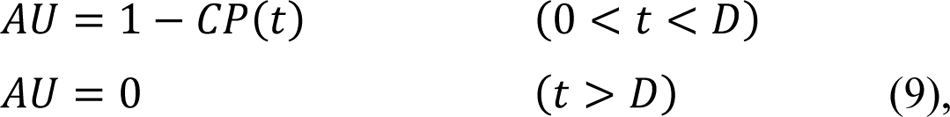

where *D* denotes the duration of the delay duration in the decision trials. Immediately after the delay finished and the reward became available, *AU* was set to 0. *AU* shows temporal dynamics that peaks at the beginning of the decision trial and monotonically decrease as the decision trial period elapses (Fig. 2D *bottom* and Fig. S2C), as in prior models (6, 22).

*CP*(*t*) is also associated with the value of a future reward that would eventually become available in the decision trial, as in prior models (6, 10, 22, 61, 62). The current study defined the future reward value as the upcoming future reward (*UFR*), formulated as *CP*(*t*) (10, 61) (Fig. S2C),

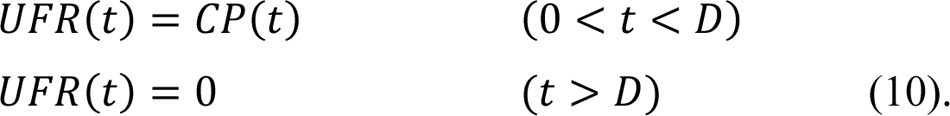

Like the *AU* value, the value of *UFR* was set to 0 immediately after the delay finished and the reward became available to drink.

The *AU* and *UFR* models were calculated for each stay trial for each participant, and then convolved with a canonical HRF (Fig. S2C *right*).

Like the survival and death functions for the leave trials, *AU* and *UFR* are anticorrelated by definition in the economic models, but when convolved with the HRF, they produced dissociable BOLD regression functions (Fig. S2D). Details about the orthogonality of the regressors and the control analyses for inspecting multicollinearity are described in the section on GLM estimations and control analyses below.

### Experience trial

For the experience trials, brain activity dynamics of anticipating a future reward were modeled similarly to the stay trials based on Bayesian inference learning. We assumed that participants 1) were unsure about when the reward would be delivered, 2) approximated the delivery time based on the previous experience trials, and 3) updated this approximation every time after participants performed an experience trial. Like the decision stay trials, the approximation was formulated by a PDF of the reward outcome using a beta distribution as a function of the expected delivery time (for details, see the “Decision trial: stay until reward delivery” section above). Unlike the stay trials, in every experience trial the participants were unable to stop waiting and had to continue to wait until reward delivery, and therefore the update was only dependent on the reward delivery time in previous experience trials. Thus, Bayesian inference was performed as

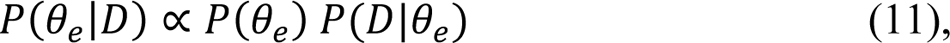

where *D* represents the delivery time of the experience trial and θ_e_ is a parameter that determines probability density. Then, *PDF* representing the approximation of the reward outcome was calculated, as in eq. (6). The average of the initial *PDF* was 15 s (i.e., average of delay durations in the practice experience trials). The *CP, AU*, and *UFR* were calculated as eqs. (7)-(10). The *AU* and *UFR* models were calculated for each experience trial for each participant, and then convolved with a canonical HRF (Fig. 4A and Fig. S6A).

As with the stay trials, *AU* and *UFR* are anticorrelated by definition in the economic models, but when convolved with the HRF, they produced dissociable BOLD regression functions (Fig. S2D). Details about the orthogonality of the regressors and the control analyses for inspecting multicollinearity are described in the section on GLM estimations and control analyses below.

### GLM estimations and control analyses

The regressors of the survival and death models for the leave trials, and also the *AU* and *UFR* models for the stay and experience trials, were simultaneously coded in the GLM model for each participant. Other events consisted of the presentation of visual messages notifying the participant of the start of the trial start and of reward delivery, and the consumption of liquid rewards, but were not of interest to the current study. These nuisance events were coded simultaneously and separately, and convolved with a canonical HRF. Then, parameters were estimated for each voxel across the whole brain.

As stated above, the survival and death models were anticorrelated, and likewise, *AU* and *UFR* models were anticorrelated in the stay and experience trials. However, when convolved with the HRF, they produced dissociable BOLD regression functions (Figs. S2A/D and 6A). Specifically, the correlations of the regressors were 0.58 ± 0.18 between the survival and death models for the leave trials, 0.31 ± 0.06 between the *AU* and *UFR* models for the stay trials, and 0.08 ± 0.11 between the *AU* and *UFR* models for the experience trials (mean ± SD), allowing sufficient dissociation (63).

Additionally, we performed separate control GLM analyses, where only one of the two anticorrelated models was coded in GLM. In those control analyses, the effects of the temporal dynamic models were reasonably reproduced, confirming that multicollinearity of these regressors was not an issue, consistent with our previous studies (6, 22).

In the leave and stay trials of the decision trials, the survival and *AU* dynamics to model temporal changes in brain activity show similar temporal characteristics: they peak at the beginning of the decision trials and monotonically decrease as the decision trial period elapses (Figs. 2A/D and Fig. S2). Similarly, the death and *UFR* dynamics show similar temporal characteristics that are the inverse of the survival and *AU* models. We then asked which model better fit the empirical data. To address this issue, we performed supplementary GLM analyses, where the relationships between trials and GLM models were interchanged. Specifically, anticipatory dynamics during the stay and leave trials were modeled by the survival and *AU* models, respectively. Then, parameters were estimated based on a standard GLM approach.

### Group-level analysis

Maps of parameter estimations for delay period effects in the experience and decision trials were collected from all participants and subjected to group-mean one-sample t-tests based on permutation methods (5000 permutations) implemented by *randomise* in the FSL suite. Voxel clusters were identified using a voxel-wise uncorrected threshold of P < .01, and the voxel clusters were tested for significance with a threshold of P < .05 corrected by the family-wise error rate. This non-parametric permutation procedure was validated to appropriately control the false-positive rate (64). The peaks of significant clusters were then identified and listed in tables. If multiple peaks were identified within 12 mm, the most significant peak was retained.

To examine the relationships between choice strategy and anticipatory brain activity, correlations between the *AU*Cs of survival functions and *AU* effects in the decision (stay) and experience trials were explored across the brain. For the stay trials of the decision trials, statistical significance was tested similarly to the group-mean tests across the whole brain. For the HPC, given the group-mean effect of *AU* in a middle part of the HPC (Fig. 2E), the exploration of the correlation was restricted within an anatomically defined middle HPC region using Harvard-Oxford cortical and subcortical structural atlases.

For the experience trials, as we asked whether the aPFC and HPC regions showing significant *AU* effects in the stay trials also showed the *AU* effect in the experience trials, the exploration of correlations was restricted within regions of interest (ROIs). Specifically, the HPC ROI was defined anatomically and the aPFC ROI was defined based on the group-mean effect of anticipatory activity in the decision trials (Fig. S6D).

To evaluate the fit of dynamic models to the empirical data, parameter estimates of aPFC and HPC ROIs were extracted (Fig. S5). The aPFC ROIs were defined based on the coordinates in the aPFC that showed *AU* effects identified in our previous studies (6, 22). The exact coordinates were (−30, 58, −8), and ROIs were created as spheres with radii of 6 mm centered on the peaks. The HPC ROIs were defined anatomically.

### Mixed-effects GLM

The aPFC showed the *AU* effect in the stay trials (Fig. 1E), and decreased aPFC activity preceded leaving the environment (Figs. 1B/C). We then hypothesized that the aPFC anticipatory activity would predict continuous choice behavior during the decision trials. To test this hypothesis, we performed a trial-based regression analysis based on a hierarchical mixed-effects GLM (65). The model included two levels, one within-subject trial-by-trial effects, and the other between-subject effects. The lower-level within-subject effects was modeled as:

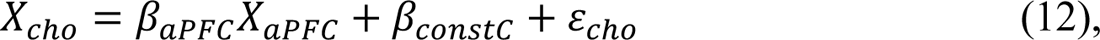

where *X_cho_* indicates (waiting duration) / (time until reward delivery during decision trials) (*X_cho_ = 1*, if participants waited until reward delivery), and *X_aPFC_* indicates the MRI signal during the decision trials in the aPFC. β-values represent regression coefficients, and ε_cho_ is an error term. The higher-level between-subject effect was modeled as:

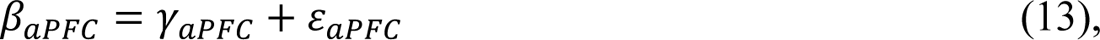

where γ-values indicate regression coefficients and *ε_aPFC_* is an error term. In this model, participants were treated as a random effect.

The aPFC ROIs were defined independently of the tested effects in order to avoid circular analysis. Specifically, the ROIs were created as spheres with a radii of 6 mm centered on the peaks based on the coordinates in our prior studies (−30, 58, −8) (6, 22).

In the ROIs, MRI signal timecourses relative to the fixation baseline were then extracted during the delay period of the decision trials. Bec*AU*se the anticipatory models (i.e., survival function and *AU*) show monotonic signal decreases toward the end of the delay period and was almost absent in the late delay period (Fig. 2A/D), the last 25% of the scanning frames of each decision trial was discarded. To eliminate a bleeding-over effect of the BOLD signal from the pre-trial period, the first four scanning frames were also discarded. Then, remaining scanning frames were averaged over the time frames for each trial. These trial-by-trial signal values were submitted to the model and all parameters were simultaneously estimated using the *lmer* procedure in R (http://www.r-project.org/). In supplementary analyses, we confirmed that fundamental results were retained when adjusting the data discard period from 0% to 50%. A separate analysis was performed by replacing the aPFC activity with HPC activity, where HPC ROI was defined anatomically.

### Functional connectivity analysis

In the stay trials, anticipatory activity was observed in the aPFC and HPC (Fig. 2E), and attenuation of aPFC activity was associated with leaving the environment (Fig. 2C).

Then, we hypothesized that the aPFC and HPC would be functionally coordinated during the stay trials where leaving did not occurred . To test this hypothesis, trial-by-trial based regression analysis was performed. Specifically, the anticipatory activity in the HPC was predicted by the anticipatory activity in the aPFC; this is formulated as

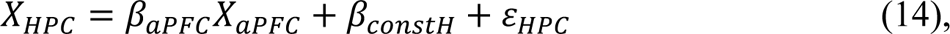

where *X_aPFC_* and *X_HPC_* indicate MRI signals during the decision trials in the aPFC and HPC, respectively. The HPC ROI was defined anatomically. MRI signals in the aPFC and HPC ROIs were calculated similarly to those used in the mixed-effects analysis above. Then, regression coefficients were estimated by the *lmer* procedure in R.

### Data and code availability

The datasets and code supporting the current study will be publicly available upon acceptance of the manuscript. (This statement will be revised accordingly).

## Supplemental information

**Fig. S1.**
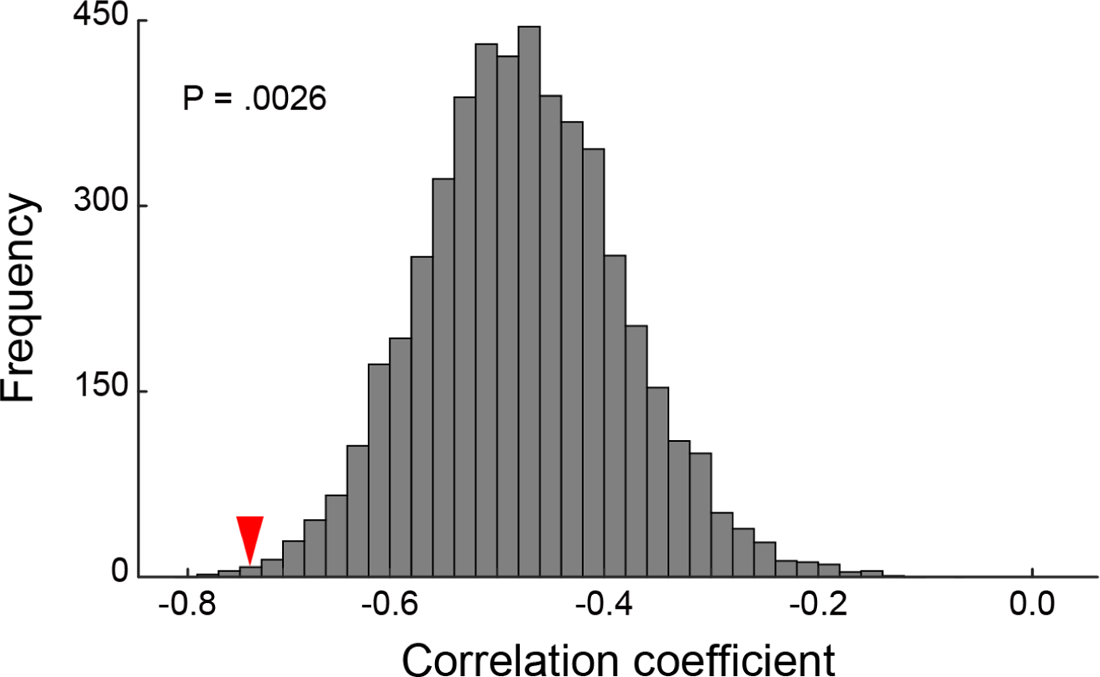
A null distribution of correlation between *AU*Cs of survival functions and sensitivity to the environmental parameters across participants. The distribution was estimated by randomly shuffling reward amount and delay duration parameters and re-labeling within individual participants. Thus, individuals’ choices were preserved when the distribution was estimated. Horizontal and vertical axes indicate the correlation coefficient, and the frequency of the permuted data, respectively. The red triangle shows the observed correlation (Fig. 1).

**Fig. S2.**
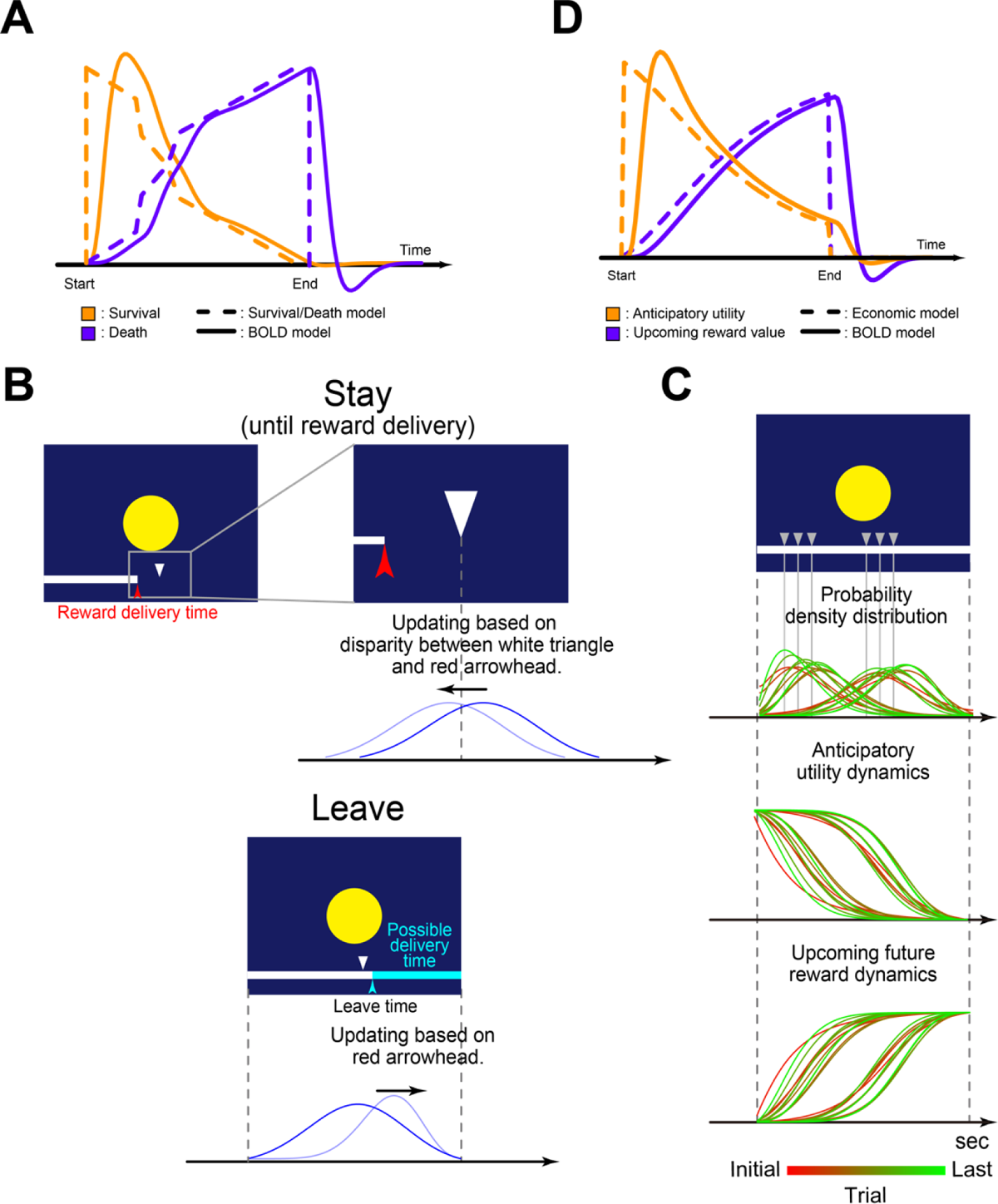
Dynamic activity models for the decision trials. A) Survival and death models for the leave trials were convolved with the canonical HRF. B) For the stay trials (*top*), the expectation of a reward outcome was updated based on 1) the reward delivery time in the experience trial (white triangle) in the environment (yellow circle) (*top left*), and 2) the disparity of the reward delivery times between the experience trial (white triangle) and the decision trial in the environment (red arrowhead) (*top right*). For the leave trials (*bottom*), the updates were based on 1) the reward delivery time in the experience trial in the environment (*bottom left*), and 2) the leave time (cyan arrowhead) that determined possible reward delivery time (i.e., somewhere between leave onset and the right end of the screen, as indicated by a horizontal cyan bar) (*bottom right*). C) Expectation of reward delivery and dynamic brain activity models. Probability density distribution for expectation of reward delivery was updated throughout the experience and decision trials (*top*). The colors of the lines indicate trial experiences as shown in the color bar at the bottom. Gray triangles indicate delivery times in the experience trials. *AU* dynamics as an inverse function of cumulative probability (*middle*). Upcoming reward value dynamics as a function of cumulative probability (*bottom*). D) *AU* and UFR models for the stay trials were convolved with the canonical HRF.

**Figure S3.**
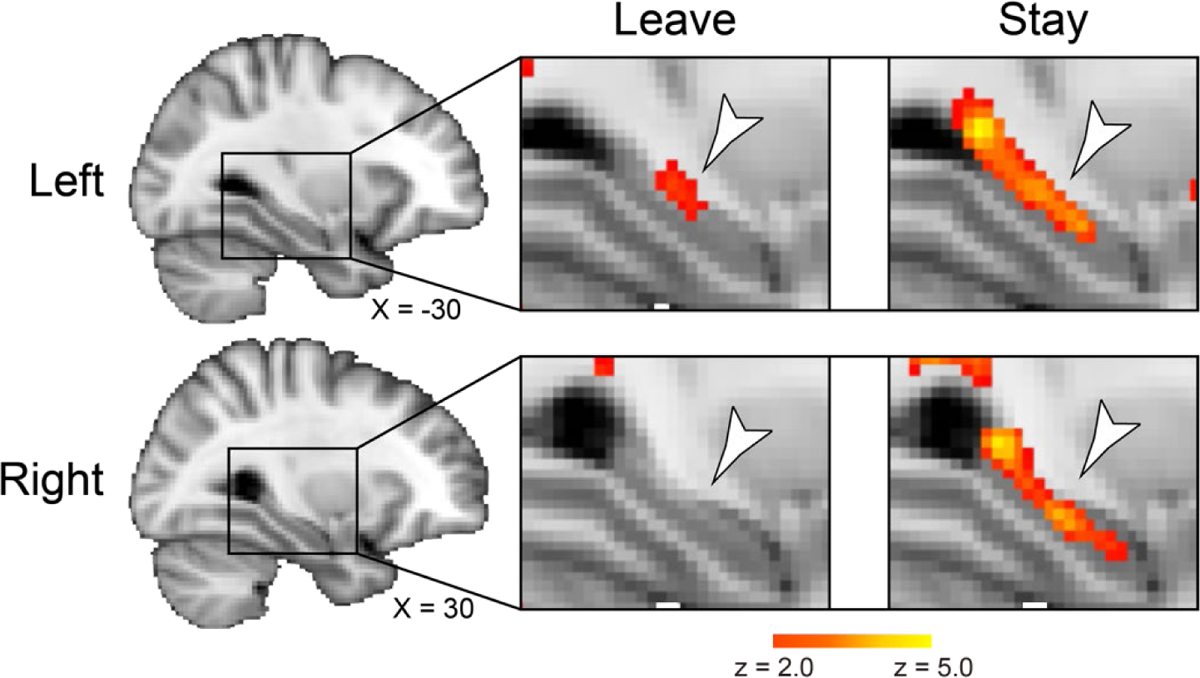
Anticipatory brain activity in the HPC (*left*) during the leave trials (*middle*) and stay trials (*right*). Statistical maps are thresholded using a voxel-wise statistical threshold of P < .05 (uncorrected) for display purposes. Participants who did not leave the current environment in any decision trial (N = 5) were excluded in the analyses. The white arrowheads indicate anatomical locations of the HPC. *Top*: left HPC; *bottom*: right HPC. Formats are similar to those in Fig. 3A.

**Figure S4.**
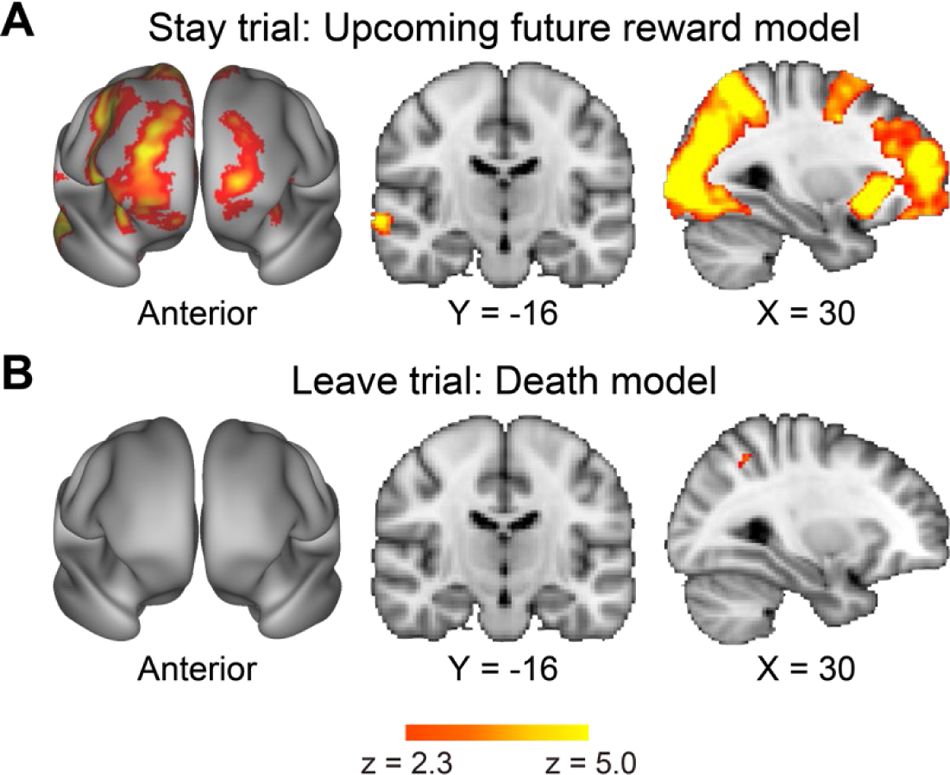
A) Brain regions showing UFR dynamics during the delay period of the stay trials. Anterior view (*left*); coronal section (*middle*); sagittal section (*right*). Levels of sections are shown below. B) Brain regions showing death model dynamics during the delay period of the leave trials. The formats are similar to the those in Fig. 2B.

**Figure S5.**
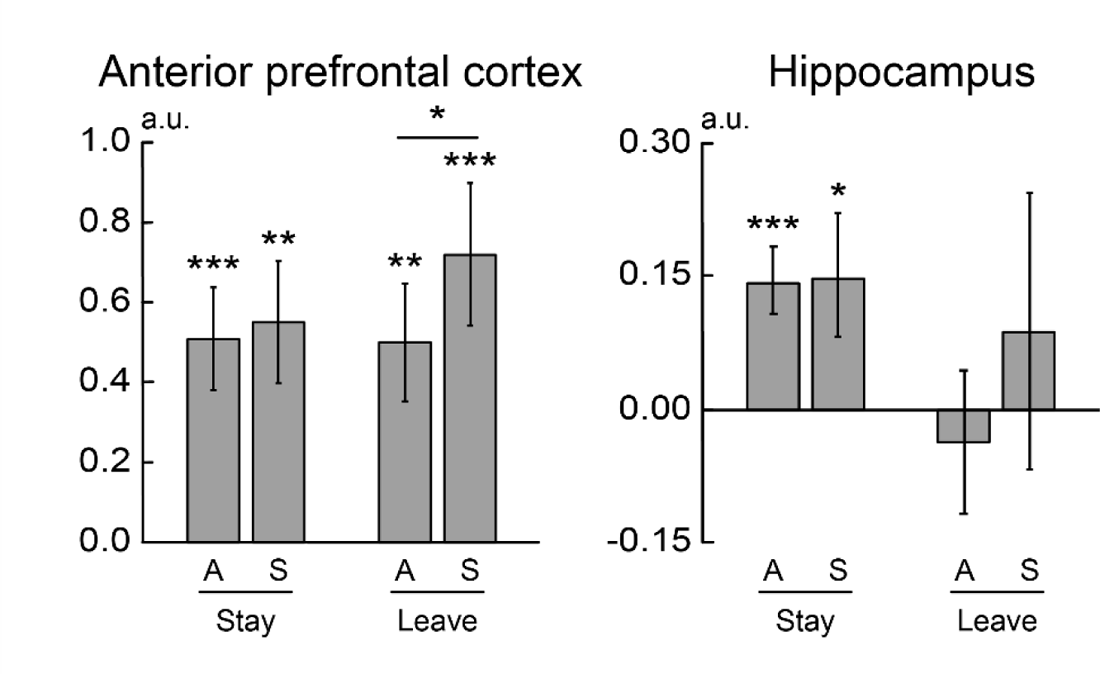
To evaluate the fit of the dynamic models for the stay and leave trials to empirical data, the relationships between trials and models (*AU* and survival) were interchanged, and parameters were estimated. Then, estimated parameters were extracted within ROIs in the aPFC and HPC. The aPFC ROI was defined by previous studies, and the HPC ROI was defined anatomically. The horizontal axis indicates trials (stay and leave) and models (A: *AU*; S: survival), and the vertical axis indicates parameter estimates. Error bars indicate standard errors of the mean across participants. *: P < .05; **: P < .01; ***: P < .001.

**Figure S6.**
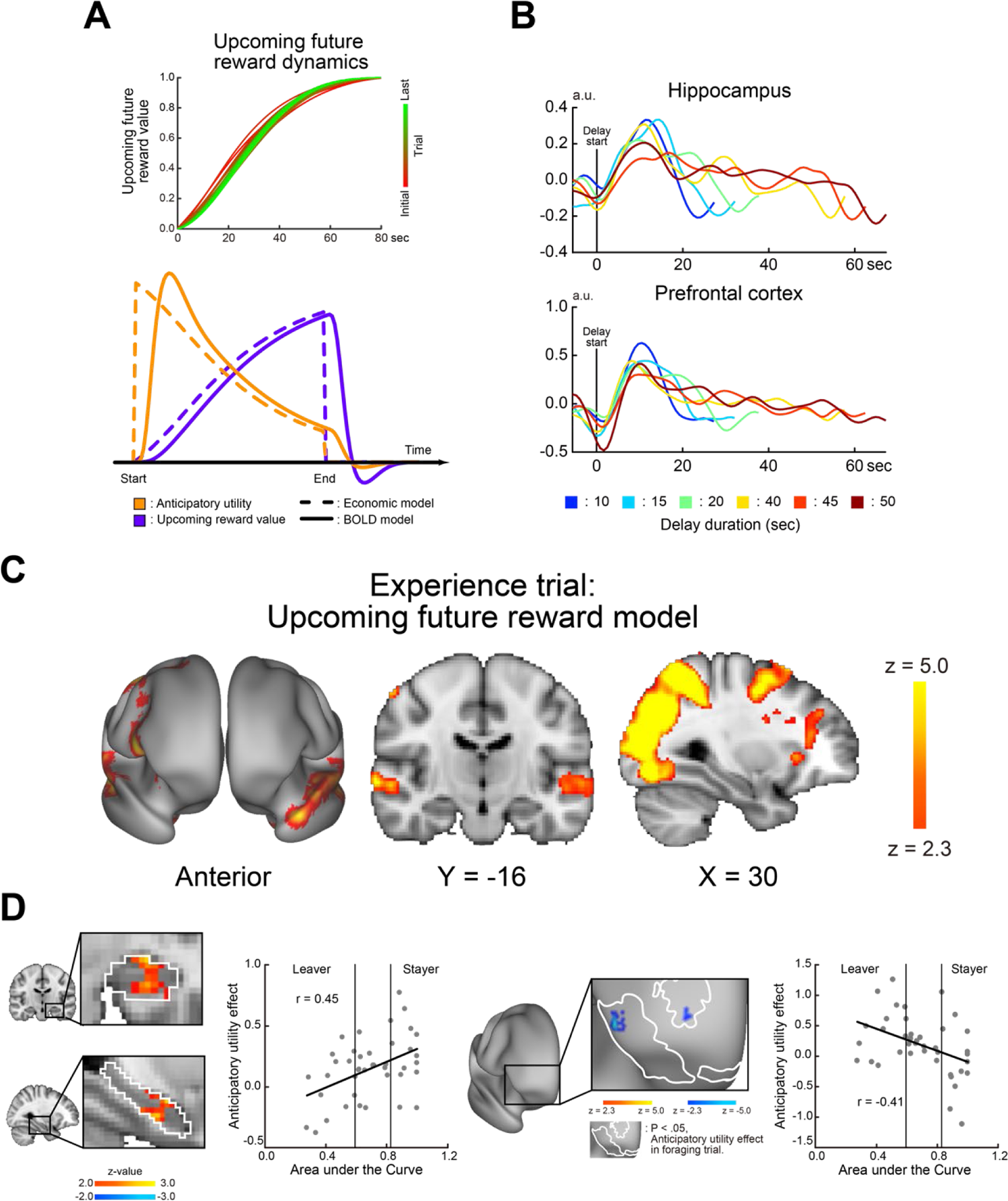
Brain activity dynamics in the experience trials. A) Dynamic models of UFR during the delay period of the experience trials (*top*). The colors of the lines indicate trial experiences as shown in the color bar on the right. *AU* and UFR models were convolved with the canonical HRF (*bottom*). B) The timecourses of activity during the delay period of the experience trials in the HPC (*top*) and prefrontal cortex (*bottom*). C) Brain regions showing UFR dynamics during the delay period of the experience trials. The formats are similar to the those in Supplementary Fig. 3A. D) Statistical maps of correlation between the *AU*C and the *AU* effect in the HPC (*left*) and prefrontal cortex (*right*). White closed lines indicate anatomical borders of the HPC and the prefrontal regions show a significant *AU* effect in decision trials in which effects in the stay and leave trials underwent weighted averaging. Formats are similar to those in Fig. 3A.

---

**Table S1.** Delay duration in the experience and decision trials (s).

| Experience |  | Decision |  |
| --- | --- | --- | --- |
| Short | 10 | 10 | 20 |
|  | 15 | 10 | 30 |
|  | 20 | 15 | 40 |
| Long | 40 | 30 | 55 |
|  | 45 | 35 | 50 |
|  | 50 | 45 | 60 |

**Table S2.**
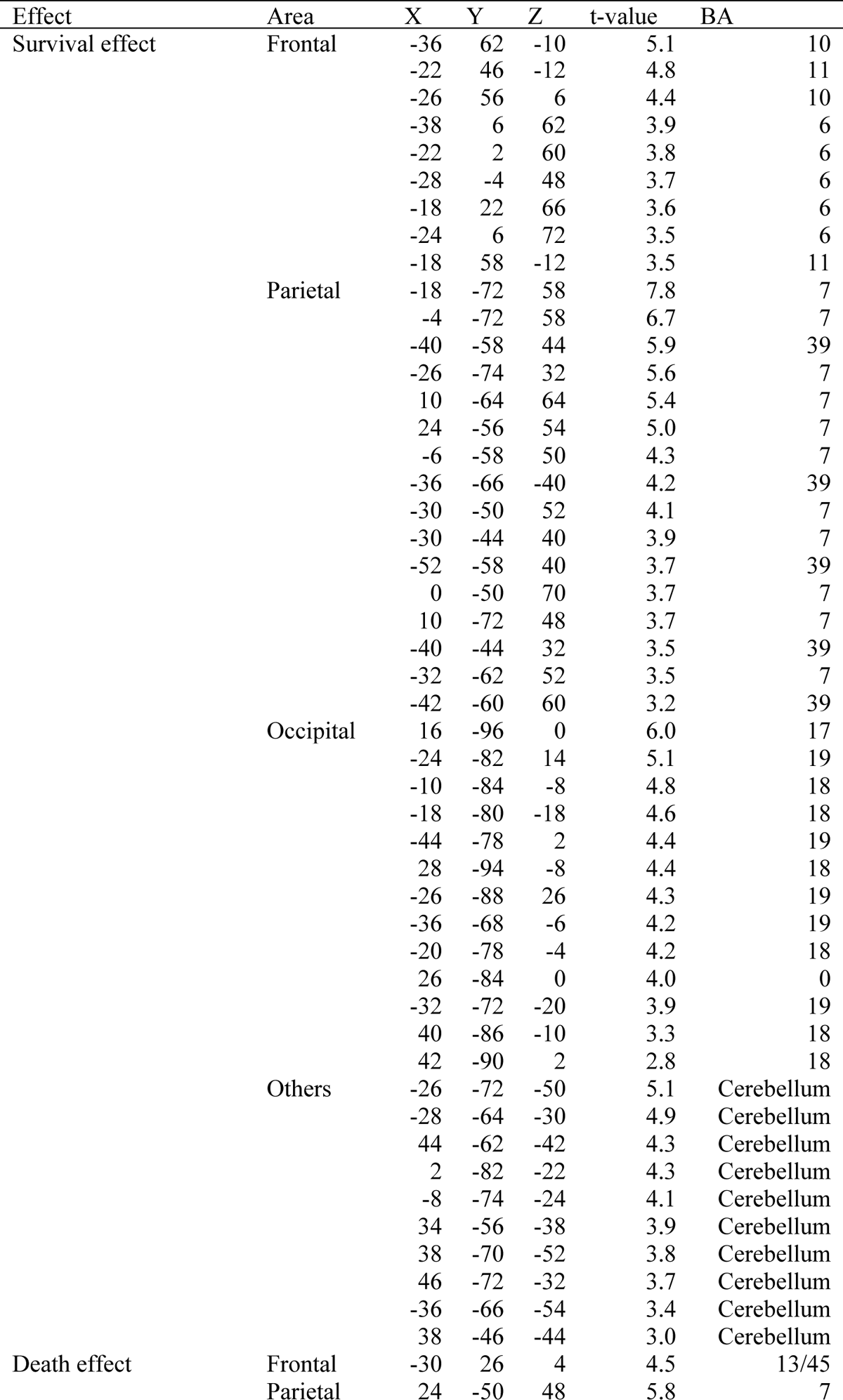
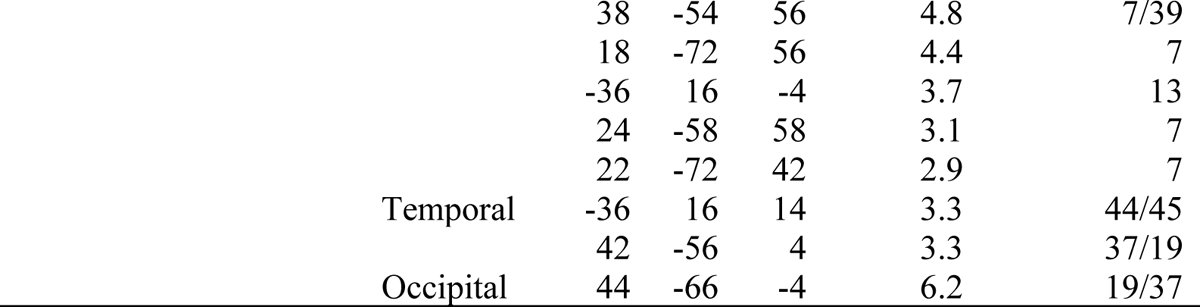
Brain regions showing significant effects of survival and death models during delay period of the leave decision trials. Coordinates are listed in MNI space. BA indicate Brodmann areas and is approximate.

**Table S3.**
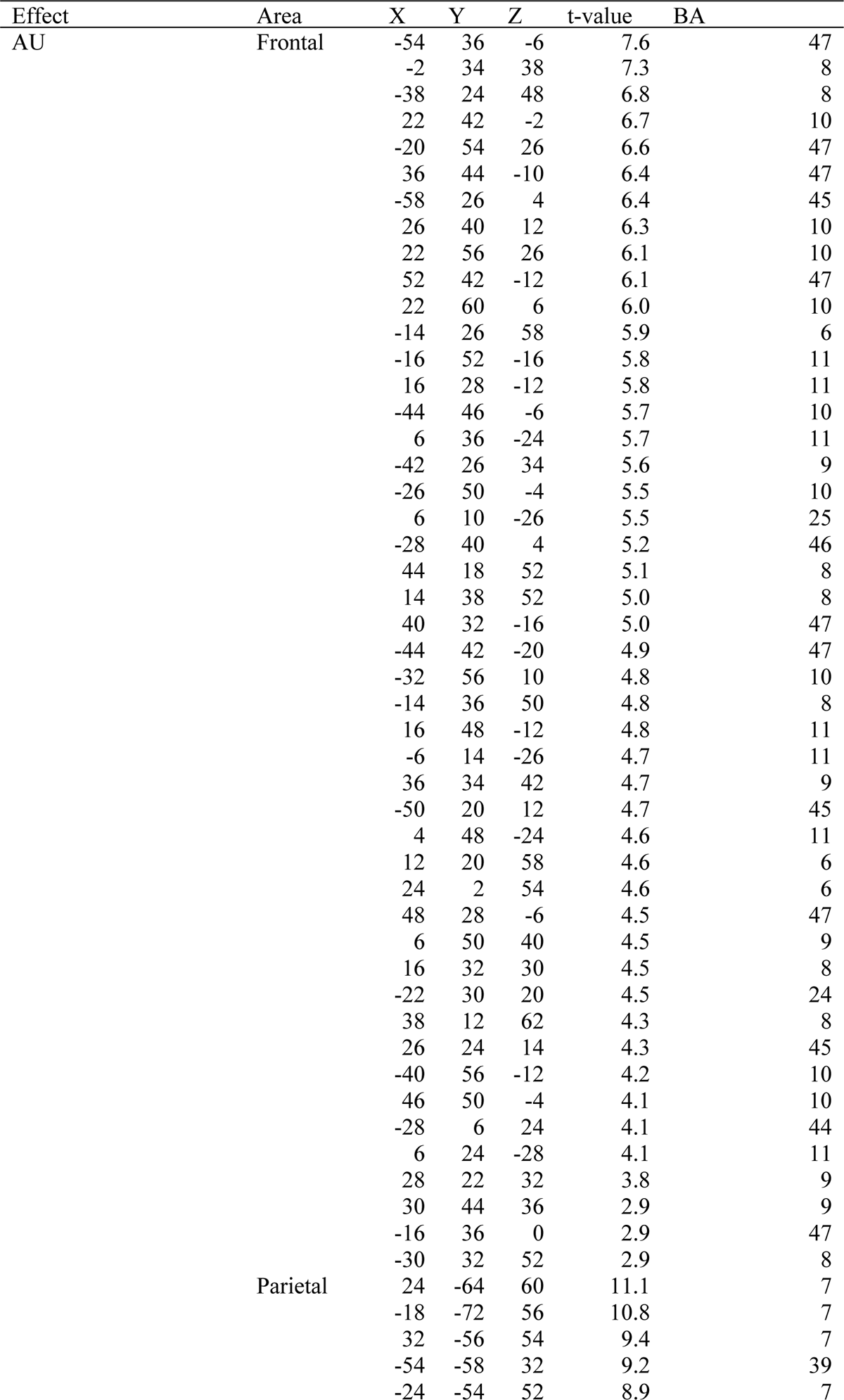
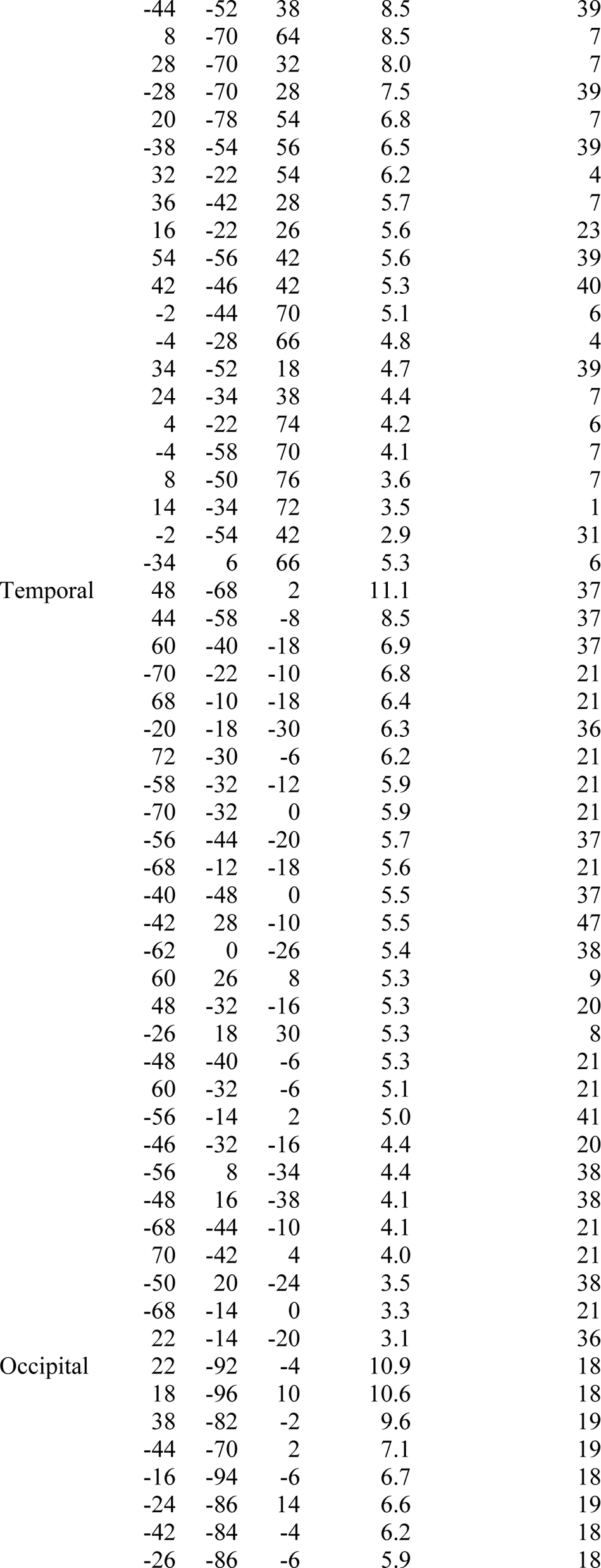
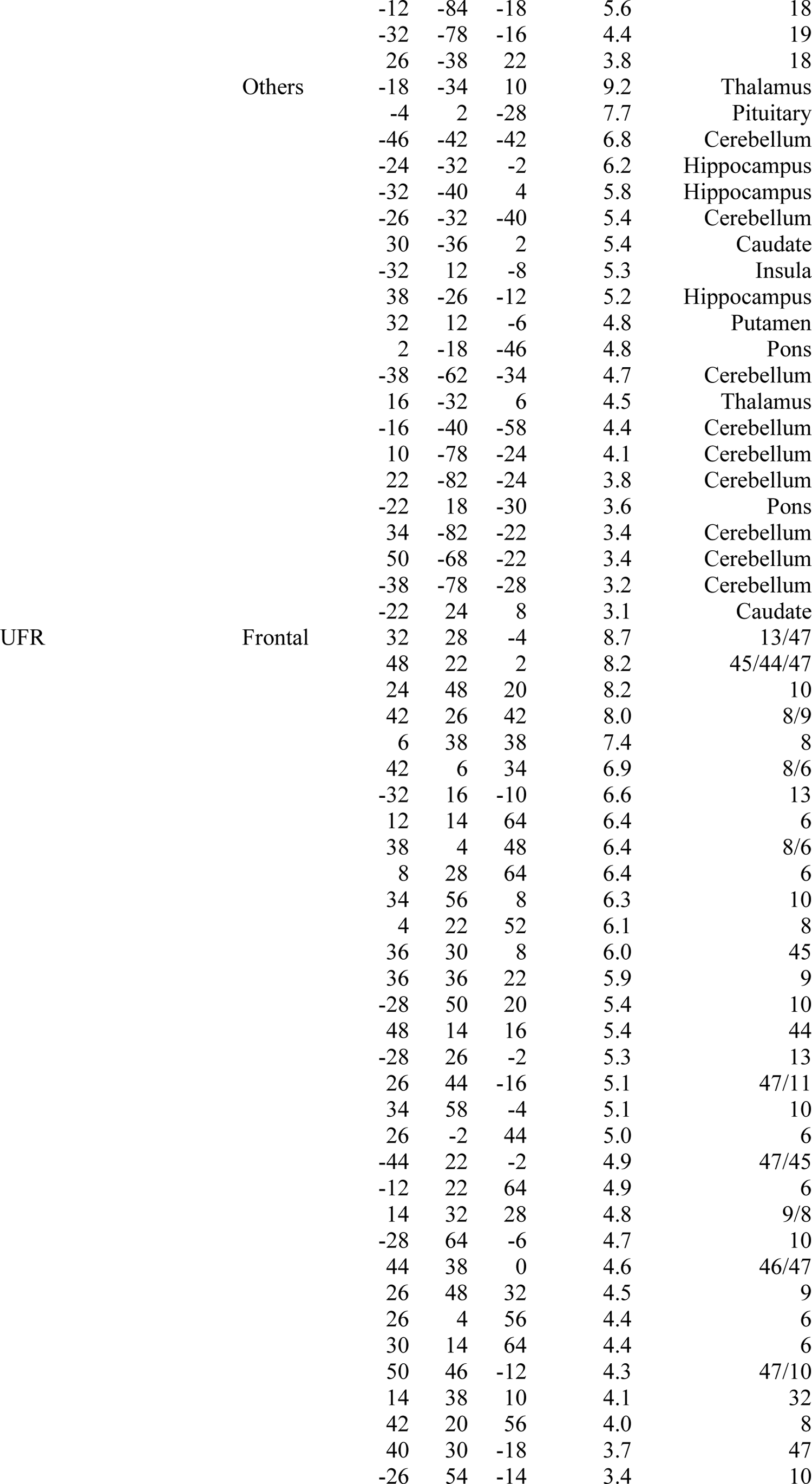
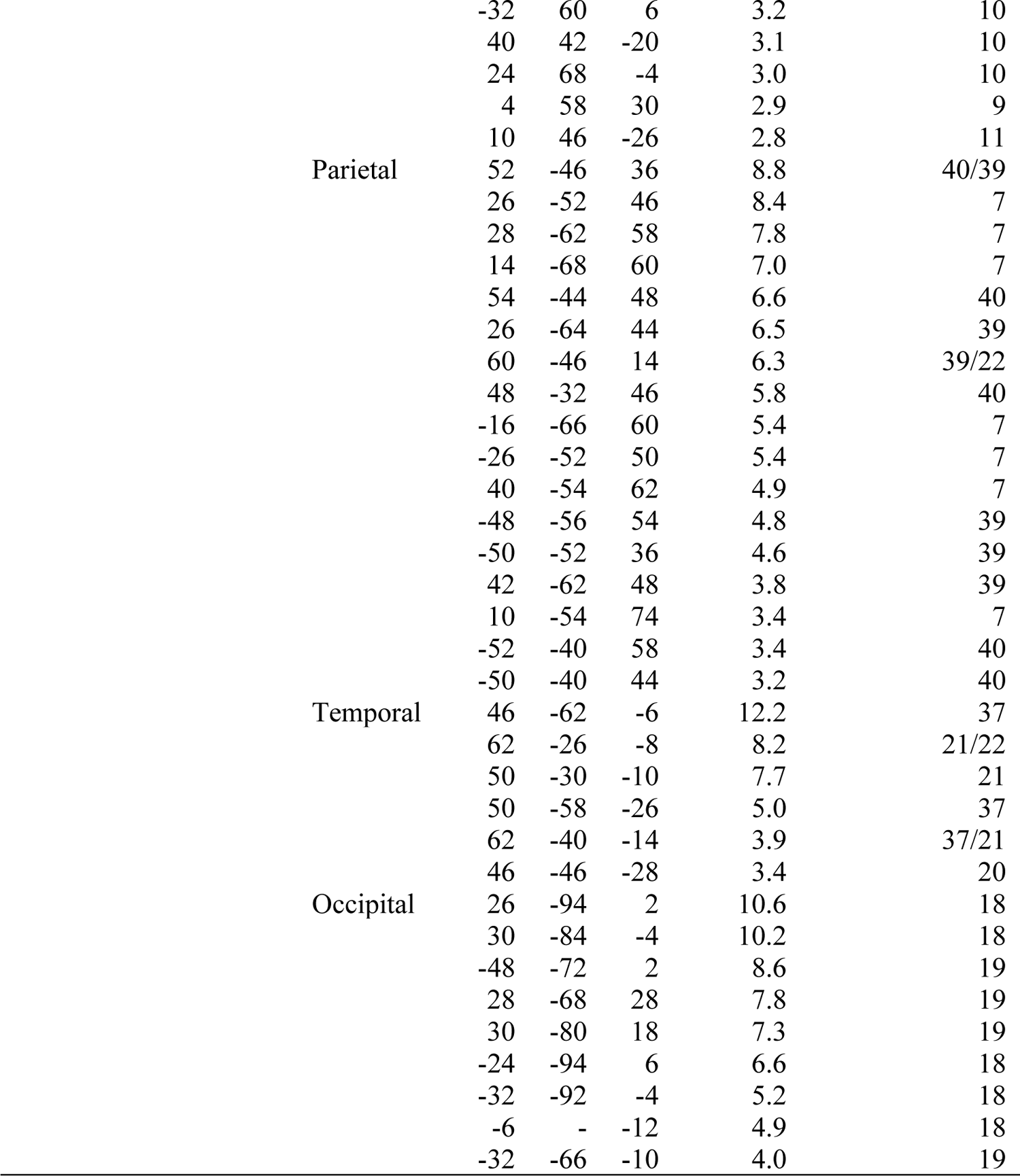
Brain regions showing significant effects of *AU* and UFR models during delay period of the stay decision trials. Formats are similar to those in Table S2.

| Effect | Area | X | Y | Z | t-value | BA |
| --- | --- | --- | --- | --- | --- | --- |
| AU | Frontal | -54 | 36 | -6 | 7.6 | 47 |
|  |  | -2 | 34 | 38 | 7.3 | 8 |
|  |  | -38 | 24 | 48 | 6.8 | 8 |
|  |  | 22 | 42 | -2 | 6.7 | 10 |
|  |  | -20 | 54 | 26 | 6.6 | 47 |
|  |  | 36 | 44 | -10 | 6.4 | 47 |
|  |  | -58 | 26 | 4 | 6.4 | 45 |
|  |  | 26 | 40 | 12 | 6.3 | 10 |
|  |  | 22 | 56 | 26 | 6.1 | 10 |
|  |  | 52 | 42 | -12 | 6.1 | 47 |
|  |  | 22 | 60 | 6 | 6.0 | 10 |
|  |  | -14 | 26 | 58 | 5.9 | 6 |
|  |  | -16 | 52 | -16 | 5.8 | 11 |
|  |  | 16 | 28 | -12 | 5.8 | 11 |
|  |  | -44 | 46 | -6 | 5.7 | 10 |
|  |  | 6 | 36 | -24 | 5.7 | 11 |
|  |  | -42 | 26 | 34 | 5.6 | 9 |
|  |  | -26 | 50 | -4 | 5.5 | 10 |
|  |  | 6 | 10 | -26 | 5.5 | 25 |
|  |  | -28 | 40 | 4 | 5.2 | 46 |
|  |  | 44 | 18 | 52 | 5.1 | 8 |
|  |  | 14 | 38 | 52 | 5.0 | 8 |
|  |  | 40 | 32 | -16 | 5.0 | 47 |
|  |  | -44 | 42 | -20 | 4.9 | 47 |
|  |  | -32 | 56 | 10 | 4.8 | 10 |
|  |  | -14 | 36 | 50 | 4.8 | 8 |
|  |  | 16 | 48 | -12 | 4.8 | 11 |
|  |  | -6 | 14 | -26 | 4.7 | 11 |
|  |  | 36 | 34 | 42 | 4.7 | 9 |
|  |  | -50 | 20 | 12 | 4.7 | 45 |
|  |  | 4 | 48 | -24 | 4.6 | 11 |
|  |  | 12 | 20 | 58 | 4.6 | 6 |
|  |  | 24 | 2 | 54 | 4.6 | 6 |
|  |  | 48 | 28 | -6 | 4.5 | 47 |
|  |  | 6 | 50 | 40 | 4.5 | 9 |
|  |  | 16 | 32 | 30 | 4.5 | 8 |
|  |  | -22 | 30 | 20 | 4.5 | 24 |
|  |  | 38 | 12 | 62 | 4.3 | 8 |
|  |  | 26 | 24 | 14 | 4.3 | 45 |
|  |  | -40 | 56 | -12 | 4.2 | 10 |
|  |  | 46 | 50 | -4 | 4.1 | 10 |
|  |  | -28 | 6 | 24 | 4.1 | 44 |
|  |  | 6 | 24 | -28 | 4.1 | 11 |
|  |  | 28 | 22 | 32 | 3.8 | 9 |
|  |  | 30 | 44 | 36 | 2.9 | 9 |
|  |  | -16 | 36 | 0 | 2.9 | 47 |
|  |  | -30 | 32 | 52 | 2.9 | 8 |
|  | Parietal | 24 | -64 | 60 | 11.1 | 7 |
|  |  | -18 | -72 | 56 | 10.8 | 7 |
|  |  | 32 | -56 | 54 | 9.4 | 7 |
|  |  | -54 | -58 | 32 | 9.2 | 39 |
|  |  | -24 | -54 | 52 | 8.9 | 7 |

|  |  |  |  |  |  |
| --- | --- | --- | --- | --- | --- |
|  | -44 | -52 | 38 | 8.5 | 39 |
|  | 8 | -70 | 64 | 8.5 | 7 |
|  | 28 | -70 | 32 | 8.0 | 7 |
|  | -28 | -70 | 28 | 7.5 | 39 |
|  | 20 | -78 | 54 | 6.8 | 7 |
|  | -38 | -54 | 56 | 6.5 | 39 |
|  | 32 | -22 | 54 | 6.2 | 4 |
|  | 36 | -42 | 28 | 5.7 | 7 |
|  | 16 | -22 | 26 | 5.6 | 23 |
|  | 54 | -56 | 42 | 5.6 | 39 |
|  | 42 | -46 | 42 | 5.3 | 40 |
|  | -2 | -44 | 70 | 5.1 | 6 |
|  | -4 | -28 | 66 | 4.8 | 4 |
|  | 34 | -52 | 18 | 4.7 | 39 |
|  | 24 | -34 | 38 | 4.4 | 7 |
|  | 4 | -22 | 74 | 4.2 | 6 |
|  | -4 | -58 | 70 | 4.1 | 7 |
|  | 8 | -50 | 76 | 3.6 | 7 |
|  | 14 | -34 | 72 | 3.5 | 1 |
|  | -2 | -54 | 42 | 2.9 | 31 |
|  | -34 | 6 | 66 | 5.3 | 6 |
| Temporal | 48 | -68 | 2 | 11.1 | 37 |
|  | 44 | -58 | -8 | 8.5 | 37 |
|  | 60 | -40 | -18 | 6.9 | 37 |
|  | -70 | -22 | -10 | 6.8 | 21 |
|  | 68 | -10 | -18 | 6.4 | 21 |
|  | -20 | -18 | -30 | 6.3 | 36 |
|  | 72 | -30 | -6 | 6.2 | 21 |
|  | -58 | -32 | -12 | 5.9 | 21 |
|  | -70 | -32 | 0 | 5.9 | 21 |
|  | -56 | -44 | -20 | 5.7 | 37 |
|  | -68 | -12 | -18 | 5.6 | 21 |
|  | -40 | -48 | 0 | 5.5 | 37 |
|  | -42 | 28 | -10 | 5.5 | 47 |
|  | -62 | 0 | -26 | 5.4 | 38 |
|  | 60 | 26 | 8 | 5.3 | 9 |
|  | 48 | -32 | -16 | 5.3 | 20 |
|  | -26 | 18 | 30 | 5.3 | 8 |
|  | -48 | -40 | -6 | 5.3 | 21 |
|  | 60 | -32 | -6 | 5.1 | 21 |
|  | -56 | -14 | 2 | 5.0 | 41 |
|  | -46 | -32 | -16 | 4.4 | 20 |
|  | -56 | 8 | -34 | 4.4 | 38 |
|  | -48 | 16 | -38 | 4.1 | 38 |
|  | -68 | -44 | -10 | 4.1 | 21 |
|  | 70 | -42 | 4 | 4.0 | 21 |
|  | -50 | 20 | -24 | 3.5 | 38 |
|  | -68 | -14 | 0 | 3.3 | 21 |
|  | 22 | -14 | -20 | 3.1 | 36 |
| Occipital | 22 | -92 | -4 | 10.9 | 18 |
|  | 18 | -96 | 10 | 10.6 | 18 |
|  | 38 | -82 | -2 | 9.6 | 19 |
|  | -44 | -70 | 2 | 7.1 | 19 |
|  | -16 | -94 | -6 | 6.7 | 18 |
|  | -24 | -86 | 14 | 6.6 | 19 |
|  | -42 | -84 | -4 | 6.2 | 18 |
|  | -26 | -86 | -6 | 5.9 | 18 |
|  |  | -12 | -84 | -18 | 5.6 | 18 |
|  |  | -32 | -78 | -16 | 4.4 | 19 |
|  |  | 26 | -38 | 22 | 3.8 | 18 |
|  | Others | -18 | -34 | 10 | 9.2 | Thalamus |
|  |  | -4 | 2 | -28 | 7.7 | Pituitary |
|  |  | -46 | -42 | -42 | 6.8 | Cerebellum |
|  |  | -24 | -32 | -2 | 6.2 | Hippocampus |
|  |  | -32 | -40 | 4 | 5.8 | Hippocampus |
|  |  | -26 | -32 | -40 | 5.4 | Cerebellum |
|  |  | 30 | -36 | 2 | 5.4 | Caudate |
|  |  | -32 | 12 | -8 | 5.3 | Insula |
|  |  | 38 | -26 | -12 | 5.2 | Hippocampus |
|  |  | 32 | 12 | -6 | 4.8 | Putamen |
|  |  | 2 | -18 | -46 | 4.8 | Pons |
|  |  | -38 | -62 | -34 | 4.7 | Cerebellum |
|  |  | 16 | -32 | 6 | 4.5 | Thalamus |
|  |  | -16 | -40 | -58 | 4.4 | Cerebellum |
|  |  | 10 | -78 | -24 | 4.1 | Cerebellum |
|  |  | 22 | -82 | -24 | 3.8 | Cerebellum |
|  |  | -22 | 18 | -30 | 3.6 | Pons |
|  |  | 34 | -82 | -22 | 3.4 | Cerebellum |
|  |  | 50 | -68 | -22 | 3.4 | Cerebellum |
|  |  | -38 | -78 | -28 | 3.2 | Cerebellum |
|  |  | -22 | 24 | 8 | 3.1 | Caudate |
| UFR | Frontal | 32 | 28 | -4 | 8.7 | 13/47 |
|  |  | 48 | 22 | 2 | 8.2 | 45/44/47 |
|  |  | 24 | 48 | 20 | 8.2 | 10 |
|  |  | 42 | 26 | 42 | 8.0 | 8/9 |
|  |  | 6 | 38 | 38 | 7.4 | 8 |
|  |  | 42 | 6 | 34 | 6.9 | 8/6 |
|  |  | -32 | 16 | -10 | 6.6 | 13 |
|  |  | 12 | 14 | 64 | 6.4 | 6 |
|  |  | 38 | 4 | 48 | 6.4 | 8/6 |
|  |  | 8 | 28 | 64 | 6.4 | 6 |
|  |  | 34 | 56 | 8 | 6.3 | 10 |
|  |  | 4 | 22 | 52 | 6.1 | 8 |
|  |  | 36 | 30 | 8 | 6.0 | 45 |
|  |  | 36 | 36 | 22 | 5.9 | 9 |
|  |  | -28 | 50 | 20 | 5.4 | 10 |
|  |  | 48 | 14 | 16 | 5.4 | 44 |
|  |  | -28 | 26 | -2 | 5.3 | 13 |
|  |  | 26 | 44 | -16 | 5.1 | 47/11 |
|  |  | 34 | 58 | -4 | 5.1 | 10 |
|  |  | 26 | -2 | 44 | 5.0 | 6 |
|  |  | -44 | 22 | -2 | 4.9 | 47/45 |
|  |  | -12 | 22 | 64 | 4.9 | 6 |
|  |  | 14 | 32 | 28 | 4.8 | 9/8 |
|  |  | -28 | 64 | -6 | 4.7 | 10 |
|  |  | 44 | 38 | 0 | 4.6 | 46/47 |
|  |  | 26 | 48 | 32 | 4.5 | 9 |
|  |  | 26 | 4 | 56 | 4.4 | 6 |
|  |  | 30 | 14 | 64 | 4.4 | 6 |
|  |  | 50 | 46 | -12 | 4.3 | 47/10 |
|  |  | 14 | 38 | 10 | 4.1 | 32 |
|  |  | 42 | 20 | 56 | 4.0 | 8 |
|  |  | 40 | 30 | -18 | 3.7 | 47 |
|  |  | -26 | 54 | -14 | 3.4 | 10 |

|  |  |  |  |  |  |
| --- | --- | --- | --- | --- | --- |
|  | -32 | 60 | 6 | 3.2 | 10 |
|  | 40 | 42 | -20 | 3.1 | 10 |
|  | 24 | 68 | -4 | 3.0 | 10 |
|  | 4 | 58 | 30 | 2.9 | 9 |
|  | 10 | 46 | -26 | 2.8 | 11 |
| Parietal | 52 | -46 | 36 | 8.8 | 40/39 |
|  | 26 | -52 | 46 | 8.4 | 7 |
|  | 28 | -62 | 58 | 7.8 | 7 |
|  | 14 | -68 | 60 | 7.0 | 7 |
|  | 54 | -44 | 48 | 6.6 | 40 |
|  | 26 | -64 | 44 | 6.5 | 39 |
|  | 60 | -46 | 14 | 6.3 | 39/22 |
|  | 48 | -32 | 46 | 5.8 | 40 |
|  | -16 | -66 | 60 | 5.4 | 7 |
|  | -26 | -52 | 50 | 5.4 | 7 |
|  | 40 | -54 | 62 | 4.9 | 7 |
|  | -48 | -56 | 54 | 4.8 | 39 |
|  | -50 | -52 | 36 | 4.6 | 39 |
|  | 42 | -62 | 48 | 3.8 | 39 |
|  | 10 | -54 | 74 | 3.4 | 7 |
|  | -52 | -40 | 58 | 3.4 | 40 |
|  | -50 | -40 | 44 | 3.2 | 40 |
| Temporal | 46 | -62 | -6 | 12.2 | 37 |
|  | 62 | -26 | -8 | 8.2 | 21/22 |
|  | 50 | -30 | -10 | 7.7 | 21 |
|  | 50 | -58 | -26 | 5.0 | 37 |
|  | 62 | -40 | -14 | 3.9 | 37/21 |
|  | 46 | -46 | -28 | 3.4 | 20 |
| Occipital | 26 | -94 | 2 | 10.6 | 18 |
|  | 30 | -84 | -4 | 10.2 | 18 |
|  | -48 | -72 | 2 | 8.6 | 19 |
|  | 28 | -68 | 28 | 7.8 | 19 |
|  | 30 | -80 | 18 | 7.3 | 19 |
|  | -24 | -94 | 6 | 6.6 | 18 |
|  | -32 | -92 | -4 | 5.2 | 18 |
|  | -6 | - | -12 | 4.9 | 18 |
|  | -32 | -66 | -10 | 4.0 | 19 |

**Table S4.**
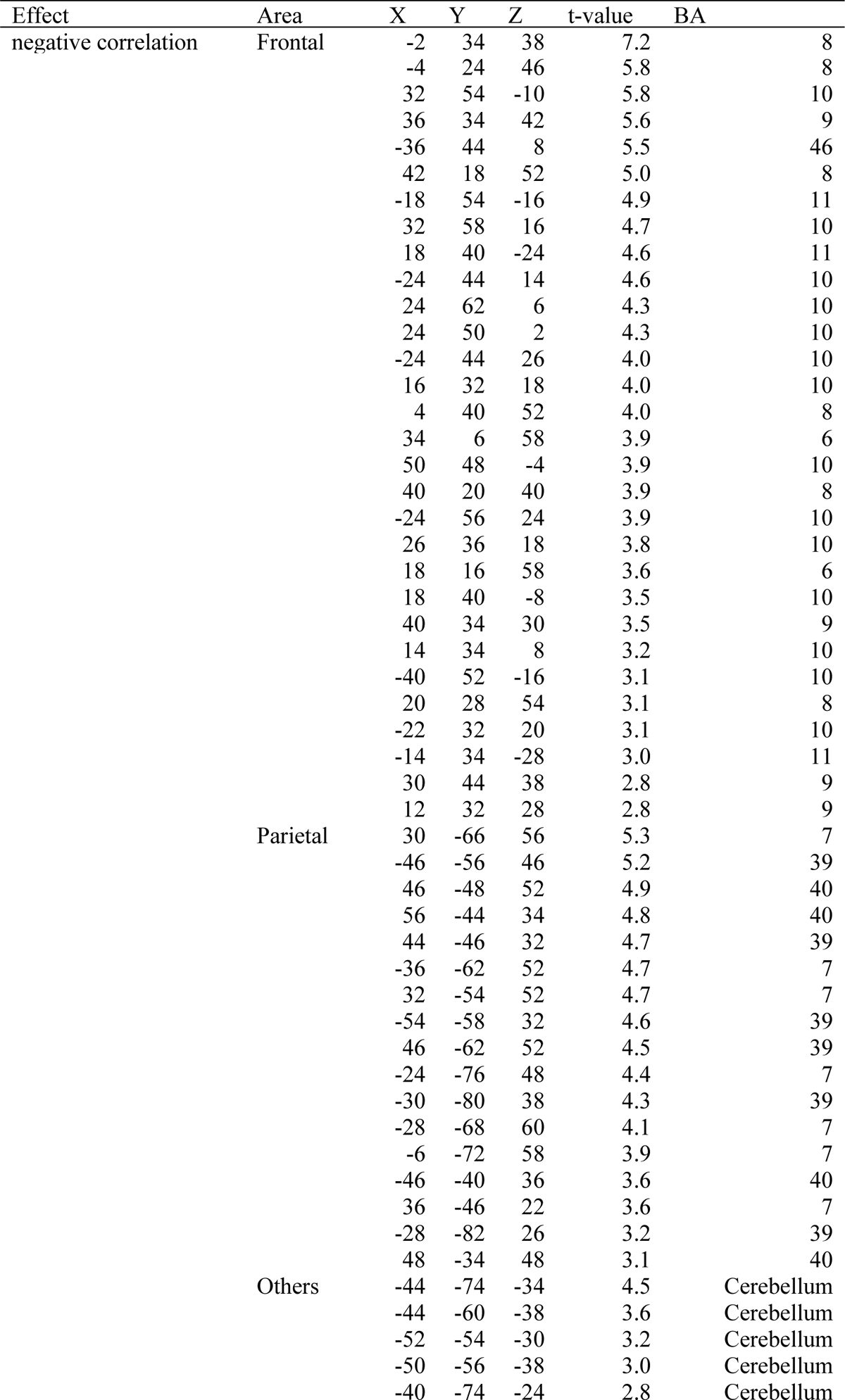
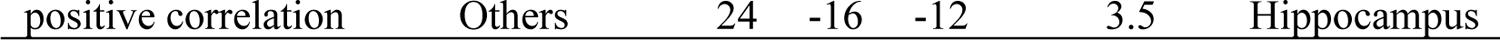
Brain regions showing significant correlations between *AU*Cs and *AU* effects during delay period of the stay decision trials. Formats are similar to those in Table S2.

**Table S5.**
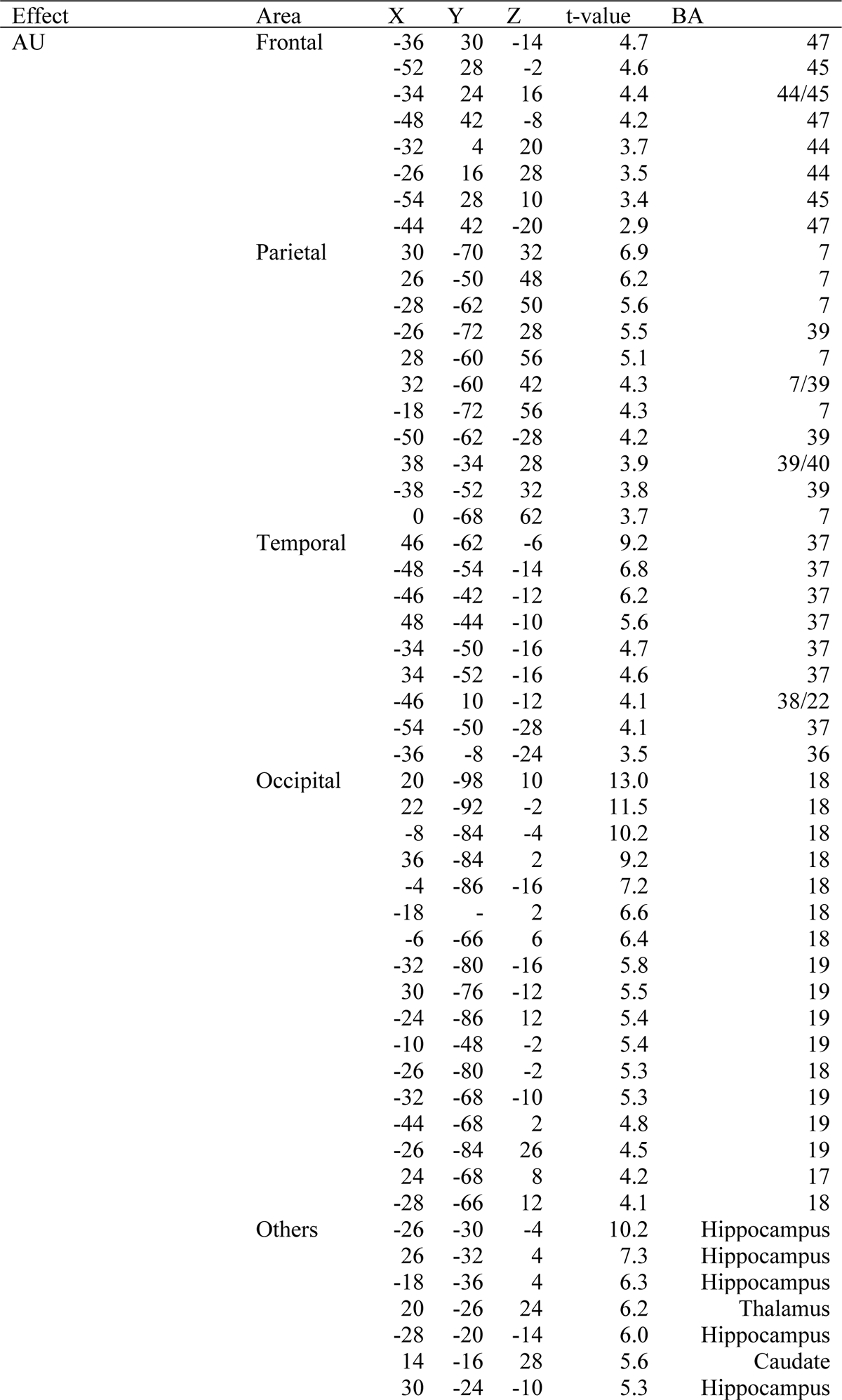
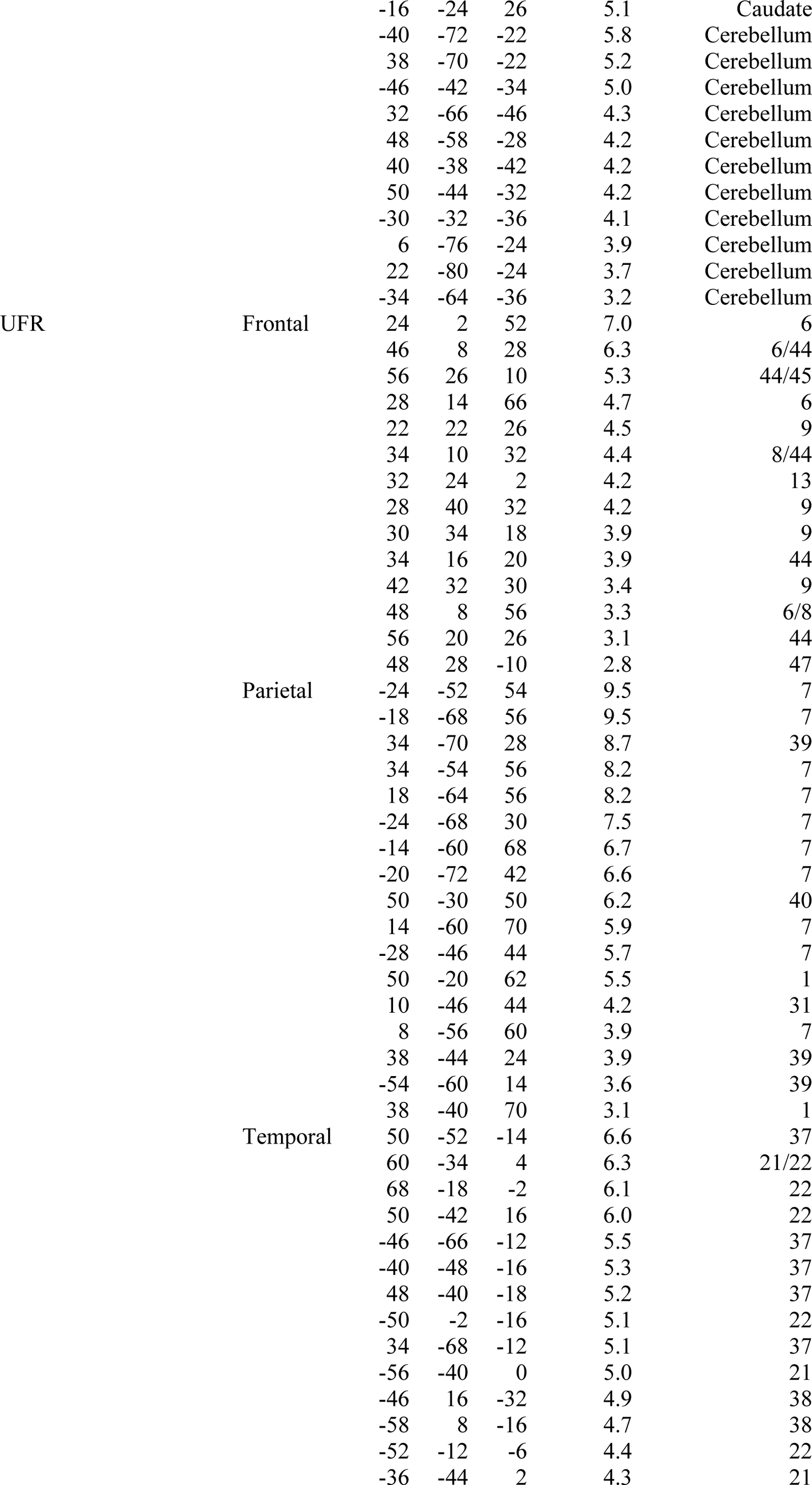
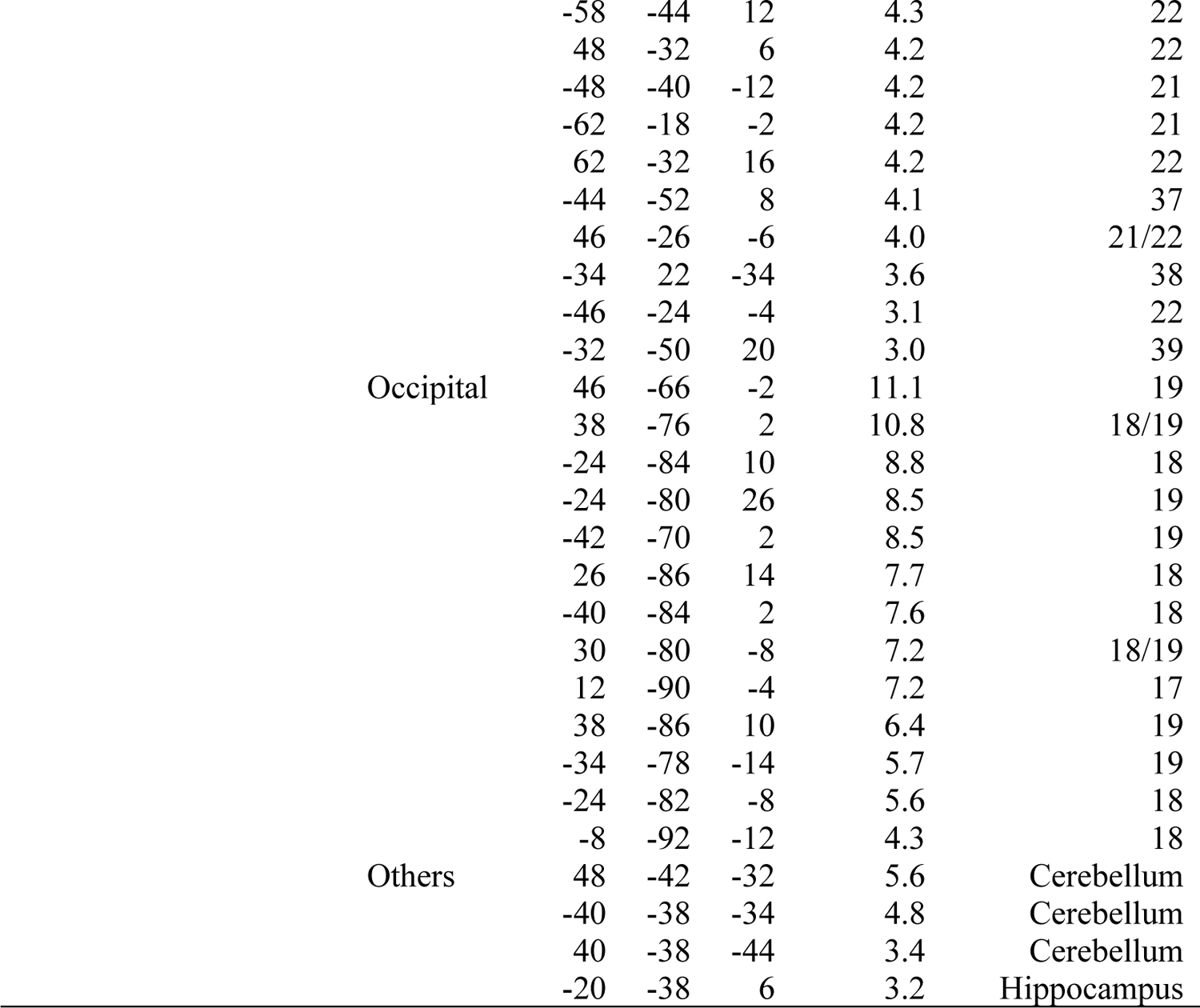
Brain regions showing significant effects of *AU* and UFR models during delay period of the experience trials. Formats are similar to those in Table S2.

**Table S6.** Brain regions showing significant correlation between *AU*Cs and *AU* effects during delay period of the experience trials. Formats are similar to those in Table S2.

| Effect | Area | X | Y | Z | t-value | BA |
| --- | --- | --- | --- | --- | --- | --- |
| Positive correlation | Others | 28 | -14 | -14 | 3.3 | Hippocampus |
| Negative correlation | Frontal | -24 | 44 | 18 | 6.0 | 10 |
|  |  | -30 | 34 | 10 | 4.1 | 45/46 |
|  |  | 26 | 56 | -6 | 3.6 | 10 |
|  |  | 40 | 48 | -4 | 3.0 | 10 |

